# Deep Learning–Based Control of Electrically Evoked Activity in Human Visual Cortex

**DOI:** 10.1101/2025.09.24.678361

**Authors:** Pehuén Moure, Jacob Granley, Fabrizio Grani, Leili Soo, Antonio Lozano, Rocio López-Peco, Adrián Villamarin-Ortiz, Cristina Soto-Sanchez, Shih-Chii Liu, Michael Beyeler, Eduardo Fernández

## Abstract

Visual cortical prostheses offer a promising path to sight restoration, but current systems elicit crude, variable percepts and rely on manual electrode-by-electrode calibration that does not scale. This work introduces an automated data-driven neural control method for a visual neuroprosthesis using a deep learning framework to generate optimal multi-electrode stimulation patterns that evoke targeted neural responses. Using a 96-channel Utah electrode array implanted in the occipital cortex of a blind participant, we trained a deep neural network to predict single-trial evoked responses. The network was used in two complementary control strategies: a learned inverse network for real-time stimulation synthesis and a gradient-based optimizer for precise targeting of desired neural responses. Both approaches significantly outperformed conventional methods in controlling neural activity, required lower stimulation currents, and adapted stimulation parameters to resting state data, reliably evoking more stable percepts. Crucially, recorded neural responses better predicted perceptual outcomes than stimulation parameters alone, underscoring the value of our neural population control framework. This work demonstrates the feasibility of data-driven neural control in a human implant and offers a foundation for next-generation, model-driven neuroprosthetic systems, capable of enhancing sensory restoration across a range of clinical applications.

## 1 Main

Implanted neuroprostheses offer a promising route to restoring lost sensory function by electrically stimulating neurons in brain areas responsible for perception. Devices targeting vision [1–3], audition [4], and somatosensation [5, 6] are in various stages of development, yet all face a fundamental challenge: interfacing with highly non-linear networks of biological neurons whose role in perception is not fully understood. Due to the limited spatial resolution of electrical stimulation, prostheses evoke neural response patterns that are unnatural to the brain, often resulting in percepts that are artificial, distorted, or variable [7–10].

One prominent example of neuroprostheses is the visual prosthesis, which electrically stimulates neurons in the retina [11–14], optic nerve [15], lateral geniculate nucleus [16], or early visual cortex [17–20] to restore visual perception. Cortical visual prostheses, which bypass the eye and optic nerve entirely, can both stimulate and record neurons in the primary visual cortex using microelectrode arrays such as the Utah Electrode Array (UEA) [17, 21]. This bidirectional interface enables the delivery of patterned stimulation and the measurement of evoked neural responses (Figure 1A).

**Figure 1:**
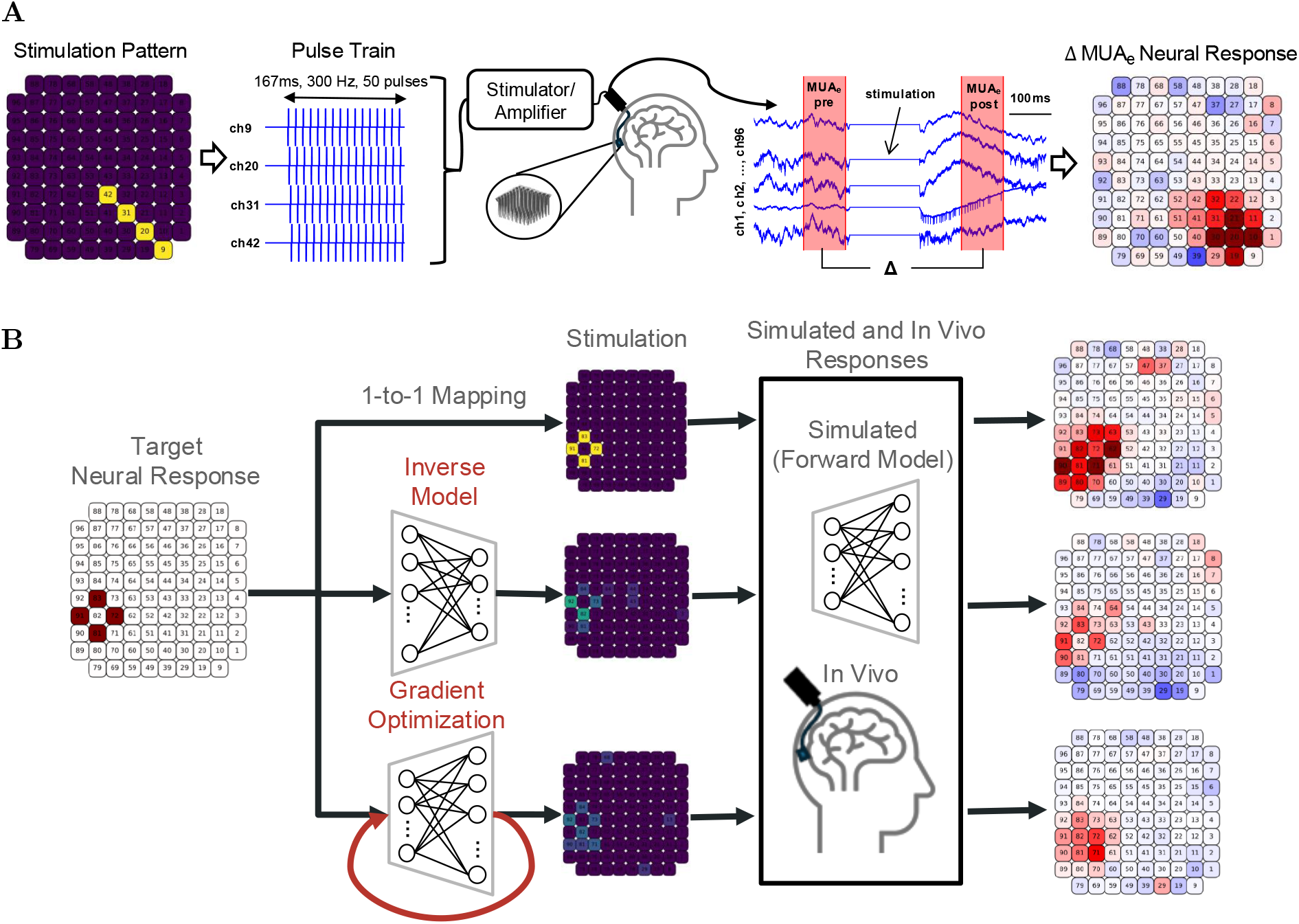
Schematic representation of the experimental setup and modeling framework. **A**) A stimulation pattern across electrodes is selected and sent to a UEA, which delivers the stimulus to the primary visual cortex of a human participant. The electrical activity under each electrode is recorded, amplified, and multi-unit spiking activity is extracted. **B**) Visualization of three approaches for neural response targeting: the conventional 1-to-1 mapping, the proposed inverse neural network method, and the proposed gradient-based optimization. Generated stimuli can then be played *in vivo* or input to the forward model to obtain real and simulated neural responses.

Despite their potential, cortical implants to date have rarely elicited percepts beyond simple shapes or flashes of light [7, 17, 18, 20]. Phosphenes (i.e., percepts induced by electrical stimulation) are often distorted, difficult to interpret, and inconsistent. Their size, brightness, shape, and color vary across time, electrodes, and individuals [17, 18]. Percepts from simultaneous multi-electrode stimulation typically do not correspond to the linear sum of individual phosphenes [17, 20], reflecting complex cortical connectivity [22, 23] and nonlinear neural integration [24]. Selecting stimulation patterns that produce meaningful and stable percepts is, therefore, nontrivial.

The conventional ‘1-to-1 mapping’ strategy assumes a direct visuotopic correspondence between visual space and the electrode grid, mapping camera pixel intensities directly to stimulation amplitudes. This ignores nonlinearities in cortical responses and perception, often resulting in distorted percepts. While alternative stimulation strategies have been proposed [25–39], none have been validated in vivo in humans using perceptually relevant pulse-train stimulation, and most do not address the substantial day-to-day variability in cortical responses that limits clinical reliability.

Recent evidence points to a tight coupling between neural responses and perception. In macaques, intracortical electrode recordings in V4 predicted the current threshold for perception [40], and cortical microstimulation could bias visual perception [41]. In humans, recorded neural activity from a UEA correlated with perceived brightness, detection thresholds, and percept duration [42, 43]. Additionally, pre-stimulus activity has been shown to influence both neural responses and perceptual thresholds [42, 44–46]. These findings suggest that closed-loop approaches, which monitor and adaptively control neural activity, could improve the consistency and quality of stimulation-evoked perception [47].

Here, we introduce a deep learning-based framework for neural activity shaping [33] in a human cortical visual prosthesis. Using multi-session recordings from a participant implanted with a UEA in early visual cortex (Figure 1A), we trained a forward neural network (NN) to predict single-trial evoked responses from stimulation parameters, incorporating pre-stimulus activity to account for day-to-day drift. This model served as an in silico testbed for two complementary control strategies: (1) a gradient-based optimizer that iteratively refines stimulation parameters to minimize response error, and (2) a learned inverse model that generates stimulation patterns from target neural responses in real time (Figure 1B). In vivo experiments demonstrate that both approaches improve control of population responses compared to 1-to-1 mapping and linear baselines, elicit more consistent percepts, and require lower currents. These findings establish the feasibility of model-driven neural control in a human visual cortical implant, representing a step toward closed-loop, scalable, and perceptually informed cortical visual prostheses.

## 2 Results

### 2.1 Deep Learning Model of Neural Responses to Electrical Stimulation

We first sought to model the relationship between electrical stimulation and evoked neural responses in human visual cortex. To this end, we collected a large dataset of stimulation-evoked activity from a 96-channel Utah Electrode Array (UEA) implanted near the primary visual cortex of a blind participant. Stimulation patterns spanned both random (N=5818, Supplementary Figure 1A) and structured (N=484, Supplementary Figure 1B) multi-electrode configurations and were delivered over the course of four months across 26 separate days. On a subset of trials (N=5515), the participant was instructed to report whether they perceived a phosphene (N=4811) and to describe it (shape, color, brightness, size; N=704). Neural responses to stimulation were recorded using the same Utah Electrode Array, from which we extracted the envelope multi-unit activity (MUA_e_). This signal reflects the pooled spiking activity of multiple neurons near each electrode tip and provides a robust estimate of local population activity [40]. To reduce variability introduced by baseline fluctuations and highlight stimulation-evoked changes, we computed a summary measure we refer to as change in envelope multi-unit spiking activity (ΔMUA_e_): the difference between the mean MUA_e_ in a post-stimulation window (100–200 ms after stimulus offset) and a pre-stimulation baseline (−110 to −10 ms). These intervals were selected to avoid contamination by stimulation artifacts while capturing short-latency evoked responses. Because responses to identical stimulation patterns varied substantially across days, we also incorporated average pre-stimulus activity from each day to account for state-dependent drift and improve across-session generalization. For more details see Methods 4.1-4.4.

The same stimulation generally evoked similar responses but with day-to-day variations (Figure 2A). For repeated stimulation patterns, the neural response’s variance across days was significantly larger than the variance within a day (Figure 2B, p*<*0.0001, Wilcoxon signed-rank test; see Methods 4.7 for details on statistical tests). We found that ΔMUA_e_ exhibited decreased variability to repeated stimuli across days than post-stimulation MUA_e_ (Figure 2C, p*<*0.001), suggesting that it is less dependent on daily short-term variations in neural state (see Supplementary Figure 3A-C). Principal Component Analysis revealed that only 48 components were required to explain 95% of the variability in ΔMUA_e_, whereas the stimulation space required all dimensions (Figure 2D). This suggests that neural responses are constrained to a lower-dimensional manifold, whereas the stimulation space is high-dimensional.

**Figure 2:**
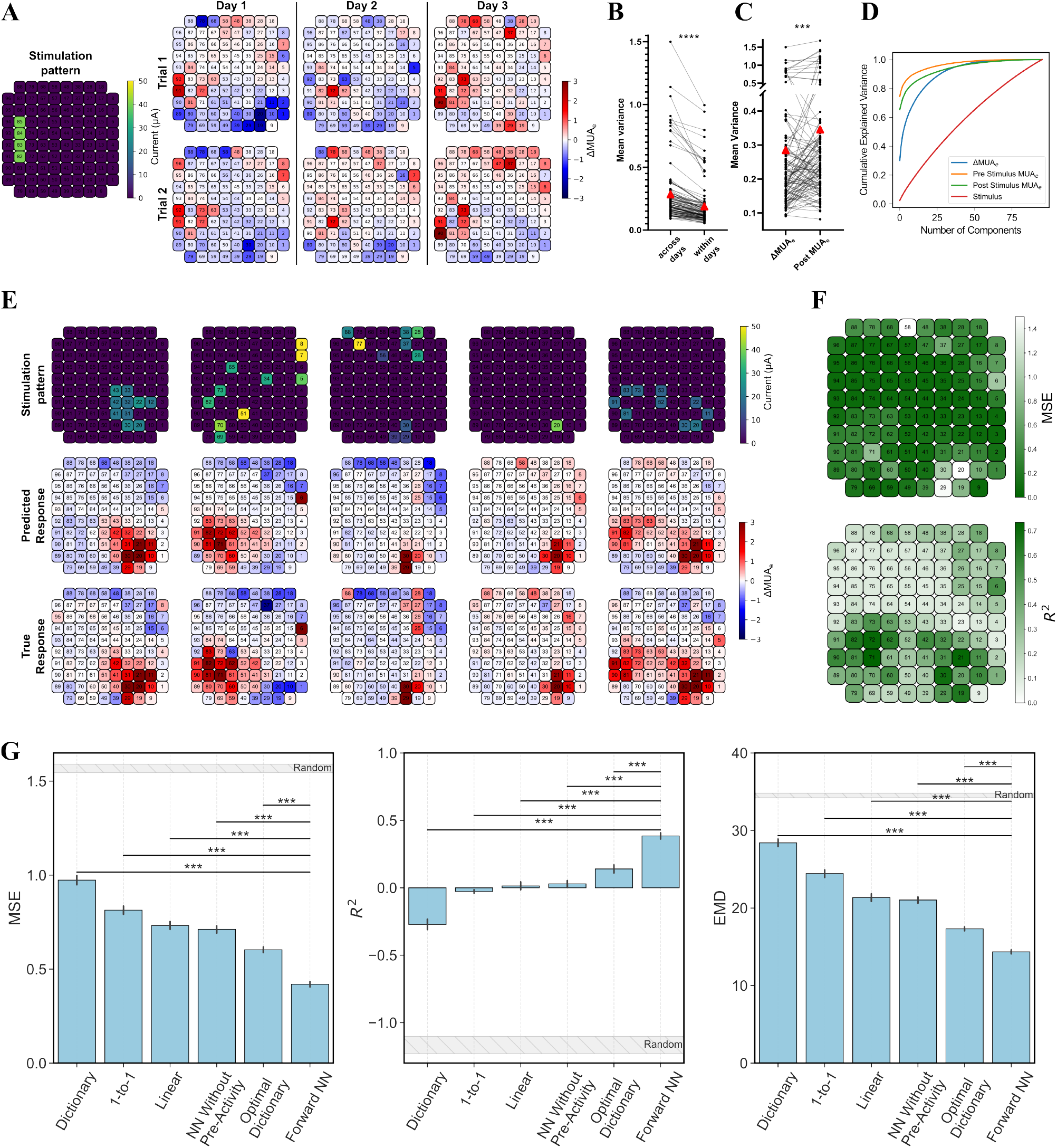
Modeling the neural response to stimulation. **A)** Example of ΔMUA_e_ over 3 days to the same stimulation pattern. **B)** Comparison of the variance of neural responses (MUA_e_) to repeated stimulations across days and within days, averaged across all channels. Red triangles represent the mean across all repeated stimulation patterns. **C)** Comparison of the variance across days of neural responses to repeated stimulations using (*left*) the difference in activity with respect to the activity before stimulation (ΔMUA_e_) and (*right*) the activity after stimulation (post MUA_e_), averaged across all channels. **D)** PCA analysis showing explained variance as a function of number of components, for stimulation patterns and neural responses, across the collected dataset. **E)** Examples of forward model predictions and in vivo neural responses on held out samples. **F)** Forward model mean squared error (MSE) and *R*^2^ for each channel. **G)** Performance of the forward model in predicting the neural response to stimulation, as measured by MSE, *R*^2^ and Earth Mover’s Distance (EMD), and compared to baseline approaches. Bars with asterisks denote significant p values: *, **, *** are *p <* 0.05, 0.01, and 0.001 respectively. Mean and standard error of the mean (SEM) shown.

From the dataset of neural responses, a deep neural network (DNN) (1.2M parameters, ‘Forward NN’, see Methods 4.8) was trained to predict the neural response (ΔMUA_e_) that would result from a given stimulation pattern. Since neural activity varied significantly day-to-day, the network was also given as input the average pre-stimulation (i.e., resting state) activity for each day. The model was trained using data from 19 days and evaluated on held-out data from 5 random non-consecutive days, resulting in a total of 5123 and 1179 train and test responses, respectively. The DNN was trained using gradient descent to minimize the MSE between the predicted and true responses.

The forward NN was compared against six baseline models (Methods 4.8): (1) a conventional receptive field model assuming focal activation and thus a one-to-one mapping between stimulation amplitude and neural activity (‘1-to-1’); (2) linear regression with access to average pre-stimulation activity (‘Linear’); (3) a dictionary-based model using an unweighted average of recorded single-electrode responses for each electrode in the multi-electrode stimulation pattern (‘Dictionary’); (4) an optimally weighted combination of single-electrode responses (‘Optimal Dictionary’); (5) the forward NN without access to pre-stimulation activity (NN Without Pre-Activity’); and (6) a random baseline using responses to random stimuli from the training set (Random’), as an estimated lower performance bound.

Performance was evaluated using MSE, *R*^2^ score averaged across electrodes, and an adapted Wasserstein EMD [48], which accounts for the spatial layout of the electrodes (Methods 4.5). Only channels which i) elicited spikes on atleast one day and ii) had a variance in neural response across all days greater than a threshold were included in the analysis, resulting in 54/96 valid channels (Methods 4.6 and Supplementary Figure 4).

The forward NN model’s predictions aligned well with recorded responses from the test set (Figure 2D). Although the MSE was low on most channels, *R*^2^ was higher on the channels near the bottom or upper right of the array, *i*.*e*. the channels with more spiking activity (Figure 2E). Our forward NN model predicted responses with significantly higher *R*^2^ and lower MSE and EMD than all baselines (Figure 2F, p *<* 0.0001). The neural network outperformed even the idealized optimal dictionary, providing strong evidence that neural responses from multi-electrode stimulation are not linear combinations of single-electrode responses.

### 2.2 Deep Neural Network-Based Control of Electrically Evoked Activity

While the forward NN enabled accurate prediction of intracortical neural responses, a critical next step is to leverage this understanding to shape neural activity. Previous attempts at electrically evoking targeted neural activity have shown varied success, often limited to simulations [33, 39], retinal explants [27], or prefrontal cortex brain states in non-human primates [49], yet never in awake humans.

To achieve this goal, two deep learning-based methods for inverting the forward neural network model were evaluated. Inspired by the success of gradient-based optimization in controlling populations of neurons via visual stimuli [50, 51], the first approach was a gradient-based method which directly searched the stimulation parameter space (*i*.*e*. the current to stimulate with on each electrode) for each target response (‘Gradient’). An iterative gradient descent algorithm optimized a stimulation pattern so that, when the stimulus was put through the forward model, the predicted neural response minimized the MSE with the target response.

Gradient descent is computationally expensive and may not be fast enough for real-time deployment. Therefore, the second NN approach consisted of training an inverse neural network (‘Inverse NN’) to map directly from target neural responses to stimuli in a single pass. This inverse network was trained alongside the forward model (with frozen weights), similar to an autoencoder [30, 33, 34]. The network learned to output stimuli that, when input through the forward model, yielded neural responses that minimized MSE with the desired target (Methods 4.9).

To evaluate these shaping approaches in vivo, optimized stimuli were generated and delivered using the prosthesis, with the participant reporting whether they perceived a phosphene. Below, we compare the performance of these approaches for both natural and synthetic target responses. Natural target responses consisted of neural activity from the test set of our collected dataset, for which there exists an original ground-truth stimulus that evoked the response on a previous day (Supplementary Figure 5). Synthetic target responses consisted of artificially crafted activity patterns with structures and shapes, potentially infeasible to evoke due to the constraints of the population of neurons under the UEA (Supplementary Figure 6).

The gradient and inverse network approaches were benchmarked against several baselines: (1) the conventional one-to-one mapping (‘1-to-1’), (2) a linear regression model (‘Linear’), and (3) an inverse dictionary approach (‘Dictionary’), where stimulation was computed as a weighted average of the previously recorded stimuli, weighted by target electrode activity (similar to a spike triggered average). For natural targets, the direct replay of stimuli that previously evoked the target responses (‘Replay Stimulus’) was included as a reference measuring day-to-day consistency in neural activity evoked by the same stimulus. A random approach was also included as a lower performance bound, where performance was averaged across responses from random stimuli in the training dataset. The neural activity shaping methods were evaluated using MSE, *R*^2^ score averaged across electrodes, and adapted Wasserstein EMD. Trials were not averaged before computing metrics. See Methods 4.9 for full details.

#### 2.2.1 Reproducing Natural Target Responses

For natural target responses, the goal was to reproduce known evoked neural responses on a new day, without knowing the original stimulation pattern. Both the inverse NN and gradient optimization significantly outperformed all baselines in reproducing the target neural responses (Figure 3A,C, p*<*.05, more examples in Supplementary Figures 7). Gradient optimization yielded the best overall performance, achieving errors comparable to replaying the original stimuli. Although gradient optimization achieved the best overall results, it required ∼10-20 seconds per example on a desktop with a NVIDIA RTX 3090. In contrast, the inverse NN’ was slightly less accurate but inferred stimuli in just ∼50 microseconds, a difference that could prove critical in real-time applications. Importantly, the perception reported by the participant for DNN-optimized stimulation more closely matched the participants perception of the original stimuli (detection task, Figure 3C).

**Figure 3:**
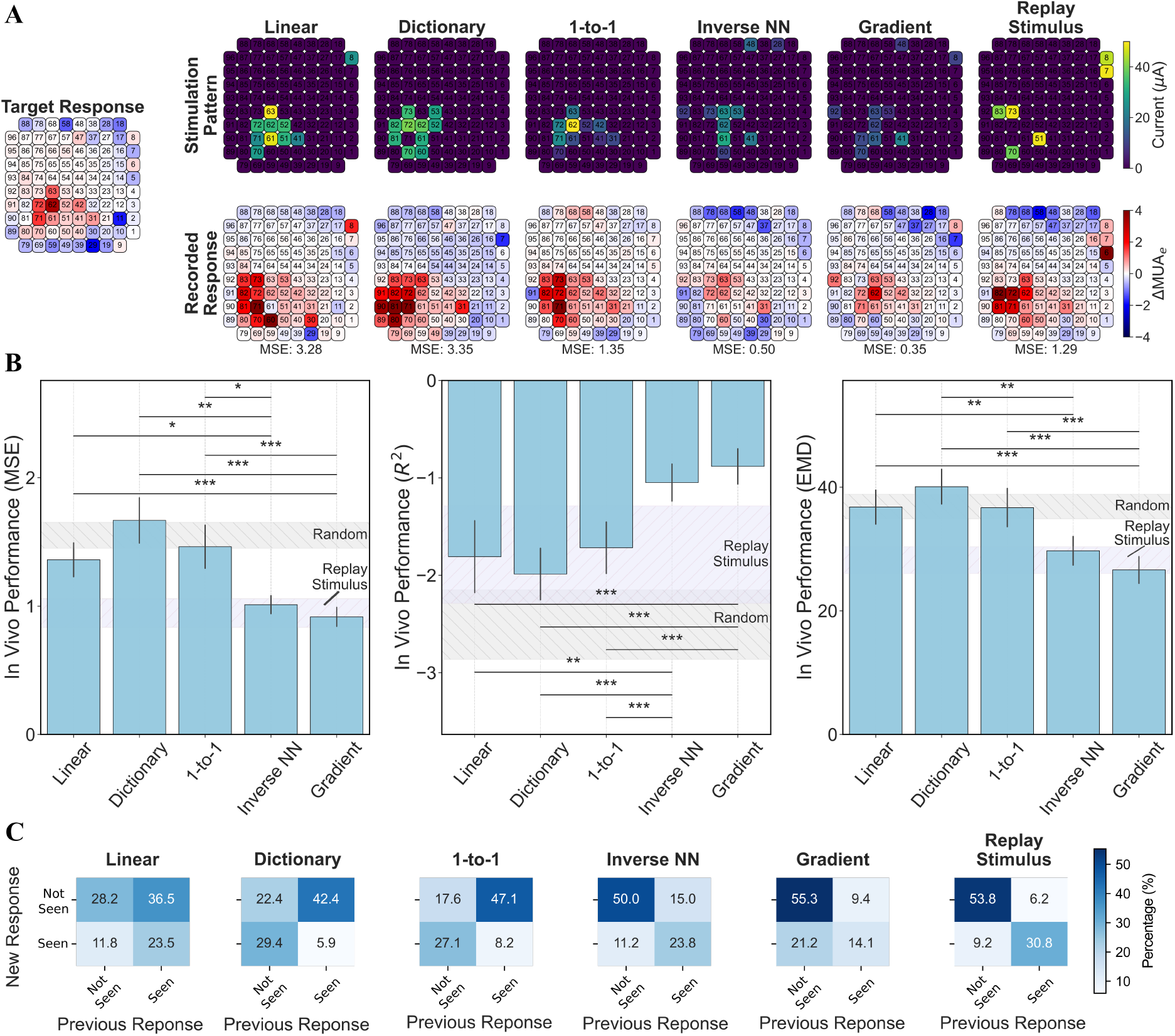
Evaluation of neural activity shaping methods. **A)** Optimized stimuli (*top*) and in vivo neural responses (*bottom*) for each shaping method for an example natural target response. **B)** Comparison of errors for each activity shaping method using MSE, *R*^2^, and EMD. Bars with asterisks denote significant p values: *, **, *** are p *<* 0.05, 0.01, and 0.001 respectively. Error bars represent SEM. **C)** Confusion matrices showing whether reported perceptions matched the previously observed perceptions associated with the same target neural activities.

#### 2.2.2 Generalizing to Synthetic Target Responses

In practical applications, we may wish to generate stimulation patterns for target responses that lie beyond the distribution of recorded neural activity. For example, real-time visual input from a glassesmounted camera could define a desired percept, but the underlying neural activity required to produce that percept may not exist within the range of previously seen responses. To test the ability of our models to generalize, we used synthetic targets: manually selected structured neural response patterns that may fall outside the feasible neural activity space (Supplementary Figure 6).

Compared to natural targets, synthetic targets had worse MSE but, interestingly, improved *R*^2^ and EMD for all methods (Figure 4). However, the relative improvement compared to random stimulation was smaller for synthetic targets than natural, suggesting that neural activity shaping was not as effective as for natural targets. Both the inverse NN and gradient approaches significantly outperformed the baseline methods in shaping neural responses (*p <*.05, Figure 4A,B, see Supplementary Figure 8 for more examples). Overall errors were higher for synthetic targets than for natural responses, suggesting constraints imposed by the stimulation paradigm, the training data, or cortical dynamics. Notably, gradient inversion again provided the closest match to the targets, while the inverse NN again showed slightly inferior performance.

**Figure 4:**
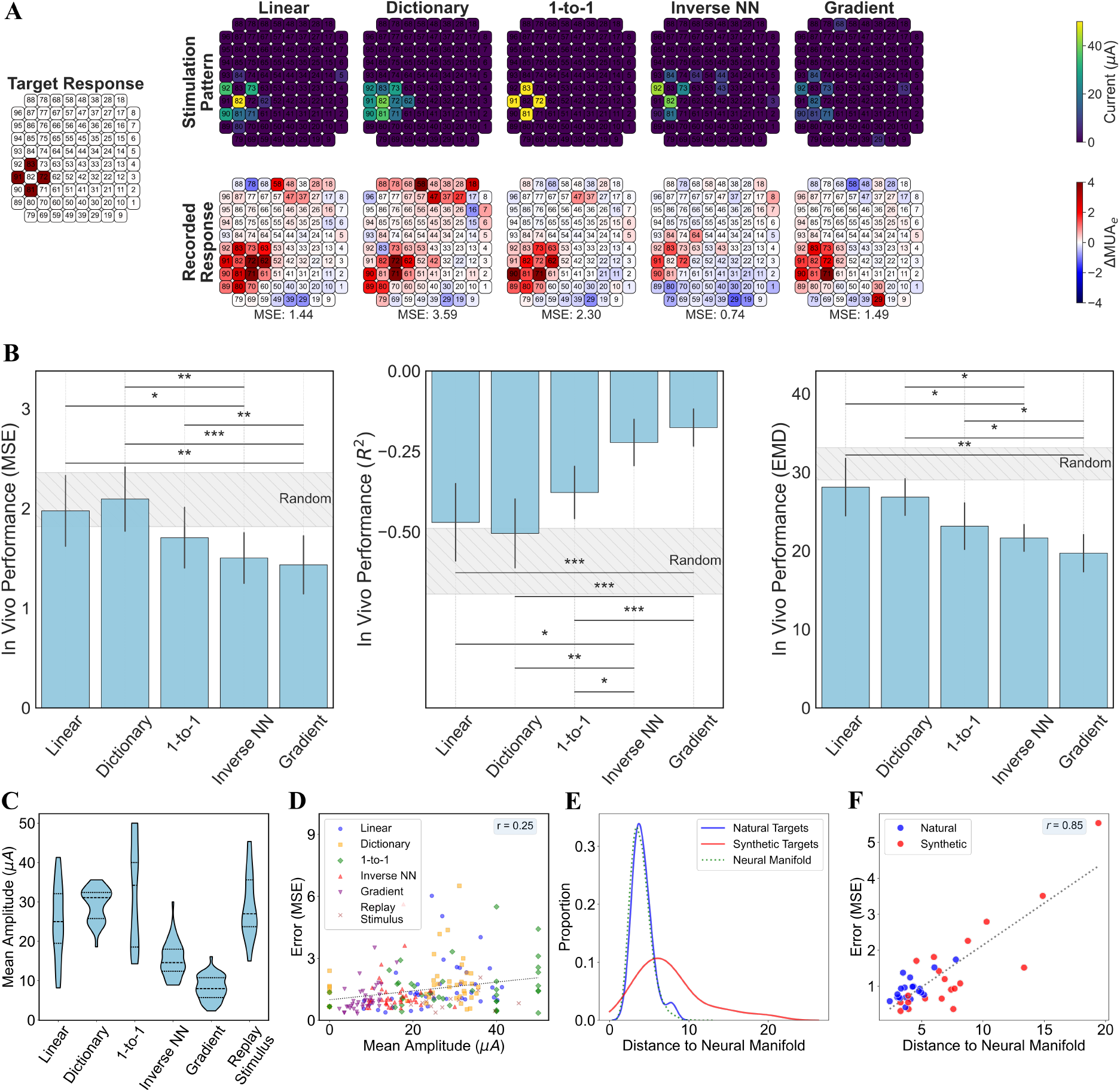
Evaluation of neural activity shaping methods for synthetic targets. **A)** Optimized stimuli (*top*) and in vivo neural responses (*bottom*) for each activity shaping method for an example synthetic target response. **B)** Comparison of errors for each method using MSE, *R*^2^, and Wasserstein EMD. Bars with asterisks denote significant p values: *, **, *** are p *<* 0.05, 0.01, and 0.001 respectively. Error bars represent SEM. **C)** Average stimulation amplitude for each method across natural and synthetic targets. **D)** Amplitude was only slightly correlated with neural response reconstruction error. **E)** Kernel density estimate plot showing the distance to the low dimensional (D=10) natural neural response manifold across target neural responses. The manifold was quantified using latent factor analysis. Synthetic targets lied further from this manifold than natural targets. **D)** Target responses which were further from the latent manifold had larger reconstruction error (shown for ‘Gradient’ responses).

Optimized stimulation patterns produced with both gradient-based optimization and the inverse NN achieved target neural responses while requiring lower overall stimulation currents (Figure 4C), representing a substantial improvement in stimulation efficiency and safety. To test whether this improvement was simply due to the use of lower amplitudes, we examined the relationship between stimulation current and model performance. Only a weak correlation was observed (*r* = 0.25, Figure 4D), indicating that the superior performance of the DNN-based approaches cannot be explained by current reduction alone.

Instead, model performance was strongly tied to a deeper constraint: the intrinsic structure of the brain’s natural activity manifold. Similar to observations in visually evoked activity [52] and other brain regions [53], electrically evoked responses were confined to a low-dimensional neural manifold, quantified using latent factor analysis [54, 55] (Methods 4.10). After accounting for noise, 95% of the manifold variance in evoked responses could be explained with just 10 latent factors, in stark contrast to the 86 factors required to explain 95% of the variance in stimulation parameters. Natural target responses clustered near the manifold, while synthetic targets deviated further (Figure 4E, *p <* 0.001, t-test). The farther a target response lay from the neural manifold, the harder it was to control, with a strong correlation between manifold distance and model error (*r* = 0.85, Figure 4F). These results suggest that precise control of evoked neural activity may be fundamentally bounded by the brain’s intrinsic response geometry, not simply by stimulation parameters.

### 2.3 Evaluating Simulated Neural Responses to Optimized Stimuli

A critical requirement for stimulus optimization is the ability to accurately predict neural responses before delivering stimulation in vivo. If the forward model can reliably approximate real-world responses, it becomes a powerful tool for designing and refining stimulation strategies without the need for extensive empirical testing. However, discrepancies between simulated and true responses must be quantified to understand the model’s limitations.

To evaluate predictive accuracy, we compared the simulated responses (i.e., those predicted by the forward model for optimized stimuli) with the true in vivo responses measured in the participant (Figure 5A,B).

**Figure 5:**
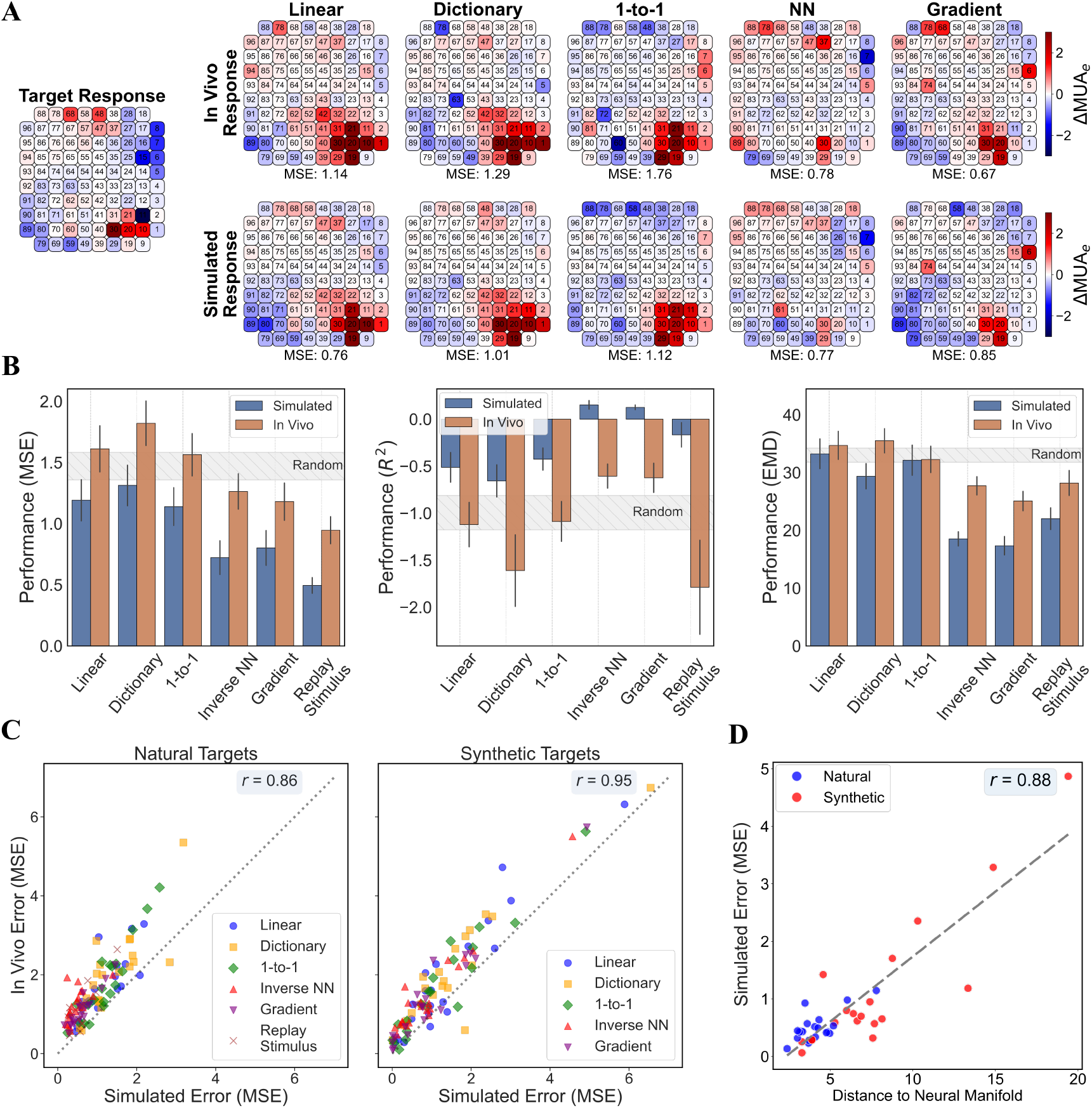
Generalization of neural activity shaping strategies from simulated models to in-vivo measurements. **A)** Simulated neural responses (*bottom*) and *in vivo* responses (*top*) for an example target. Simulated responses were obtained by running the synthesized stimulus through the forward neural network model. **B)** Comparison of errors for simulated and in vivo responses across methods. **C)** Simulated vs in vivo neural response reconstruction error (MSE) for each shaping method, for natural (*left*) and synthetic (*right*) targets. **D)** Target responses with a larger distance to the latent neural manifold also had larger simulated errors.

Across all inversion techniques, simulated responses systematically overestimated the accuracy of neural activity shaping compared to in vivo performance (lower MSE and EMD, higher *R*^2^). However, this overestimation was consistent across methods, and the relative ranking of the different stimulation synthesis techniques was roughly preserved.

Crucially, despite the tendency to overestimate performance, simulated and in vivo MSE were highly correlated across targets (*r* = 0.86 for natural and *r* = 0.95 for synthetic targets; Figure 5C). Thus, while absolute errors differed, simulation results reliably predicted which targets and strategies would perform better in vivo, making the forward model a valuable surrogate for optimization.

Further supporting this, we found that even within the simulation, the difficulty of achieving a target response was strongly tied to its distance from the neural manifold. The simulated MSE was highly correlated with the simulated target’s manifold distance (Figure 5D), reinforcing that deviations from the brain’s natural activity structure impose fundamental limits on the controllability of neural responses, both in silico and in vivo.

### 2.4 Predicting Perception from Neural Responses

A critical question in the design of stimulation strategies is whether predicting neural activity actually improves perceptual outcomes. One might argue that it would be more direct (and more useful) to predict perception from stimulation parameters alone, bypassing the intermediate step of neural response modeling. To test this, we evaluated whether recorded neural activity provides a better predictor of perception than stimulation parameters themselves.

deep neural network (DNN) decoder models were trained to predict the participant’s verbal reports of phosphene detection (seen or not seen, *N* = 4416), brightness (subjective rating 1-5, *N* = 427), and color (4 color categories, *N* = 427). Decoders were trained using different input features: stimulation parameters alone, neural responses alone (ΔMUA_e_ or ΔMUA_e_ plus average pre-stimulus MUA_e_), or combinations of these inputs. Models were trained via gradient descent to minimize classification error, with performance evaluated using leave-one-day-out cross-validation to assess generalization across recording sessions (see Section 4.11 for details).

Neural activity significantly outperformed stimulation parameters alone in predicting detection, color, and brightness ratings (Figure 6A-C, p*<*0.05). When including both pre-stimulus MUA_e_ and change in envelope multi-unit spiking activity (ΔMUA_e_) alongside stimulation parameters (a realistic scenario for future visual prostheses), accuracy improved from 74.2% to 88.7%, corresponding to 56% reduction in error. Similar improvements were observed for color prediction (49.4% to 76.7%, 54% error reduction) and brightness prediction (44.8% to 60.9%, 29% error reduction). F1 scores (Figure 6D-F) confirmed these trends.

**Figure 6:**
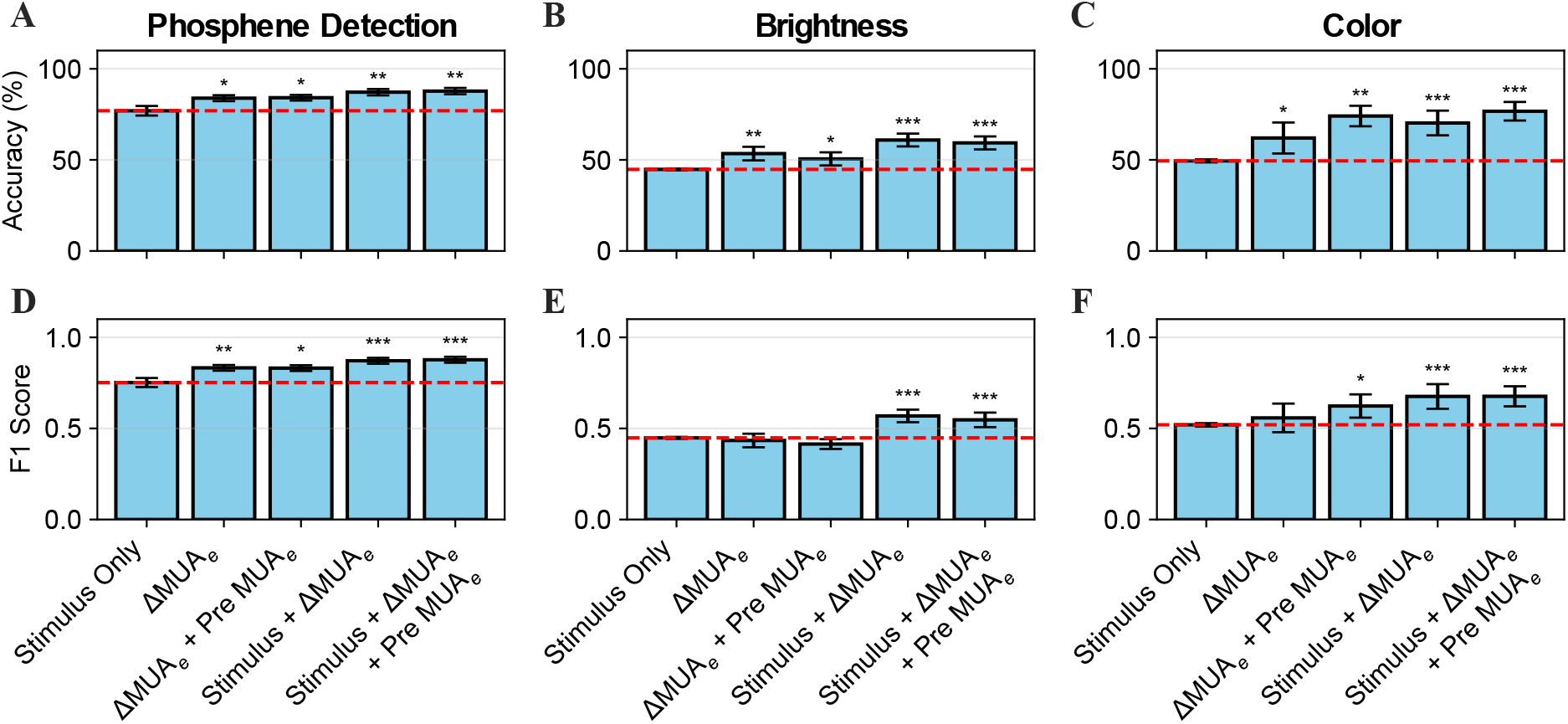
Perceptual relevance of evoked activity patterns. **A-C)** Accuracy of decoder models for phosphene detection (*N* = 4416), brightness (*N* = 427), and color (*N* = 427) perception, decoding using different input features. Leave-one-day-out cross validation was used, and performance aggregated across predictions from different folds (mean ± SEM). Asterisks denote significant differences when compared to stimulus only: *, **, *** are p *<* 0.05, 0.01, and 0.001 respectively. The dashed line is the stimulus only condition, repeated for clarity. **D-F)** F1 scores for the same perceptual attributes and model inputs as A-C.

Together, these results demonstrate that recorded neural activity provides substantially more information about perception than stimulation parameters alone. This finding supports the use of neural response modeling as a critical component of any framework aiming to optimize perceptual outcomes in cortical visual prostheses.

## 3 Discussion

This study demonstrates that deep learning-based stimulus optimization enables robust control of electrically evoked neural activity in the human visual cortex. Using a forward model trained on months of intracortical recordings, we achieved accurate prediction of neural responses across substantial day-to-day variability, allowing stimulation strategies to be refined in silico. Our deep learning-based inverse models successfully shaped neural activity to match target patterns, significantly outperforming conventional methods, while requiring lower stimulation currents and eliciting more consistent perception. Critically, we showed that neural activity, not stimulation parameters alone, provides the strongest predictor of perceptual outcomes, underscoring the central role of neural response modeling for optimizing sensory restoration. While demonstrated here in the context of cortical visual prostheses, this framework provides a generalizable foundation for adaptive, closed-loop control of brain activity across a broad range of bidirectional neural interfaces.

### Necessity of Nonlinear Neural Activity Models

When predicting neural responses, our deep NN forward model significantly outperformed multiple linear approaches (Figure 2G), providing evidence that neural responses to electrical stimulation are highly nonlinear. This presents a challenge for cortical prostheses, as many traditional stimulus optimization techniques rely on linear receptive field mapping. Our results highlight the necessity of nonlinear, scalable models that can effectively capture the complex relationship between stimulation and neural responses, particularly as implants incorporate more electrodes and higher-density arrays.

This nonlinearity was further reflected in the performance of the inverse models, where deep learning-based approaches significantly outperformed conventional stimulus optimization methods (Figure 3 and Figure 4). We evaluated two inverse models: (1) an inverse network, which maps target neural responses to stimuli in a single pass, and (2) a gradient-based optimization method, which iteratively refines stimulation parameters to minimize response error.

Both approaches achieved superior neural control compared to 1-to-1 mappings and linear models, with gradient-based optimization yielding the best performance (lower MSE, EMD, and higher *R*^2^). However, the computational cost of *∼*20 seconds per stimulus limits its current real-time feasibility. In contrast, the inverse network performed inference in *∼*50 microseconds, making it a practical choice for real-time applications. Future implementations may benefit from hybrid approaches, balancing fast inference with periodic fine-tuning to further optimize neural activity shaping.

Activity shaping had smaller performance improvements for synthetic target responses than for natural target responses. It remains an open question whether these discrepancies arise due to fundamental limitations of cortical stimulation (*e*.*g*., due to current spread, underlying neural circuitry, or activation of passing axons) or the modeling constraints imposed by generalizing outside the training distribution. If the network were trained on a broader set of response patterns, it is possible that previously unachievable targets could be brought into the distribution of feasible responses, thereby improving generalization. However, the fact that the forward model predicted out-of-distribution stimulus responses reasonably well (Figures 2 and 5), and the inverse models were trained using the forward model, suggests that at least part of the error might be due to more fundamental constraints on cortical microstimulation.

Notably, we found that electrically evoked neural responses lied on a low-dimensional manifold, and that the distance from a target response to this natural manifold was highly correlated with the error when eliciting that target. While it has been suggested that visually evoked neural activity is distributed according to a lower-dimensional manifold than the stimulus space ([52], but see also [56]), it is more surprising that the same holds for electrically evoked activity. If this is a fundamental constraint due to underlying neural circuitry, then this evoked neural manifold will have important implications for visual prostheses. Future studies will need to determine whether these limitations reflect intrinsic constraints of the neural system or whether they can be mitigated through adaptive training paradigms and expanded stimulation strategies.

### Generalization and the Forward Model as a Predictive Tool

A key finding of this study is the strong correlation between simulated and in vivo neural responses (Figure 5), validating the forward model as a predictive tool for designing new stimulation strategies. Even for synthetic targets, the model provided consistent and generalizable approximations of neural responses. The strong correlation between simulated and in vivo responses enables a computationally scalable search for optimal stimulation patterns without requiring extensive in vivo testing. This capability is particularly valuable for refining stimulation protocols and adapting prostheses for individual users.

One challenge in neurostimulation is the variability of neural responses over time, where the same stimulation pattern elicits different responses on different days. This variability likely contributes to the fluctuations in perception reported in prior studies [17]. Our approach partially mitigates this issue by considering the difference in neural activity before and after stimulation (ΔMUA_e_), which is more stable across days (Figure 2A-C). Additionally, by conditioning stimulation patterns on the average daily pre-stimulation state, our method adapts to variations in latent brain activity. Note that there was no significant performance improvement conditioning instead on per-trial pre-stimulus activity, meaning that accounting for daily drifts in activity was sufficient for shaping neural responses.

### Perceptual Relevance and Implications for Visual Prostheses

Our DNN optimized stimuli evoked perceptions more closely matching the perceptions associated with natural target responses (Figure 3C), while requiring lower stimulation amplitudes (Figure 5C). Additionally, perceptual descriptions were decoded more accurately when including neural responses than from stimulation parameters alone (Figure 6), reinforcing the idea that optimizing neural activity—rather than solely stimulation parameters—could enable more precise and reproducible perceptual outcomes.

One question that remains is determining the target neural response associated with a desired perception. Using image neural networks predictive of primate and human V1 could be one solution [39, 50], but our perception decoders might offer another solution. These decoders could be included in the training loop, allowing for perception-driven and neural activity-aware stimulus optimization. This aligns with previous work [41] demonstrating the potential of model-informed microstimulation on modulating perception in sighted primates. By integrating these perception models with our shaping strategy, we could develop fully data-driven approaches for controlling prosthetic vision that exploit the relative abundance of neural activity data, in order to optimize each individual user’s perception.

### Limitations and Future Directions

Despite these promising results, several limitations remain. First, our study was conducted in a single participant, and generalization to broader patient populations remains an open question. Second, while our model effectively captured nonlinear neural responses, it does not account for long-term plasticity or adaptive changes in cortical processing over time. Finally, although our inverse models outperformed traditional methods, further improvements are needed to enhance the real-time feasibility of gradient-based approaches.

Future work should explore multi-participant studies, investigate long-term neural plasticity in response to stimulation, and refine hybrid models that combine fast inference with adaptive optimization. Additionally, integrating closed-loop control mechanisms (i.e., where neural activity is continuously monitored and stimulation is dynamically adjusted) could significantly improve the stability and robustness of perception over time.

### Towards Closed-Loop Visual Neuroprostheses

These findings help lay the foundation for a closed-loop neural prosthesis, shifting the focus from fixed stimulation patterns to adaptive neural activity shaping guided by predictive models. This has broad implications for sensory brain-computer interfaces, particularly as bidirectional implants (which can both stimulate and record neural activity) become more widespread. By leveraging deep learning models to dynamically adjust stimulation in response to ongoing neural activity, future prostheses could achieve greater stability, precision, and perceptual fidelity in real-world applications.

## 4 Methods

### 4.1 Electrode Array and Participant

The experiments were conducted as part of a clinical research trial for the development of a cortical visual prosthesis for the blind based on intracortical microelectrodes. The clinical trial was approved by the Clinical Research Committee of the General University Hospital of Elche and registered at ClinicalTrials.gov (NCT02983370). The study adhered to the principles outlined in the Declaration of Helsinki, and all relevant ethical guidelines associated with clinical trial regulation were followed, including EU No. 536/2014 (repealing Directive 2001/20/EC) and EU Commission Directives 2005/28/EC and 2003/94/EC. The participant provided written informed consent prior to their participation. The confidentiality and privacy of the participant were maintained throughout the study, and all procedures were designed to minimize any potential risks to their well-being.

A blind male volunteer (27 years old) with good physical and mental health took part in the clinical trial. He had lost his vision due to a traumatic head injury, after which bilateral enucleation was performed one year prior to his participation.

At the start of the trial, one UEA was implanted into the early visual cortex as an interface for the cortical visual prosthesis [21, 57]. The implant was placed in the right occipital cortex, likely in or near V1 based on cortical anatomy. However, fMRI verification of its precise location was not possible due to the presence of metal fragments in the skull.

The UEA consists of 100 1.5-mm-long electrode shanks (96 of which effectively connected to the pedestal connector) and can both electrically stimulate the visual cortex to induce visual perception and record the neural activity. A robot-assisted procedure was followed in the UEA implantation [58] in addition to the standard surgical procedure [17]. A platinum wire implanted under the dura was used as reference electrode. After 6 months, the subject underwent another surgery where the UEA was explanted.

During the 6 months of implantation (after which the subject was explanted), the subject performed daily sessions where electrical stimulation of the visual cortex induced visual perceptions (phosphenes). Experimental sessions usually lasted 4 hours per day, 5 days per week. The data collected for this work span across 26 daily sessions, with the last 2 sessions dedicated to the optimized stimulation patterns. The visual perceptions he described were dots or half-circle shapes, and varied in size, color and brightness. Single electrode thresholds were found on 27/96 electrodes. Thresholds ranged from 5 to 90 *µ*A (mean 24 *µ*A, standard deviation 18 *µ*A).

### 4.2 Neural Data Recording and Electrical Stimulation

The Ripple Neuromed Summit Explorer processor was used for both neural recording and electrical stimulation. It was connected to the UEA connector through three Micro2+Stim front-ends (32 channels each). Neural signals were sampled at 30 kHz from 96 channels applying a 0.1 Hz – 7.5 kHz analog filter. The resolution of the analog-to-digital conversion was set to 0.25 *µ*V (16 bit). The stimulation step-size was set to 1 *µ*A. The trigger of electrical stimulation was recorded as a digital event in the stimulation data stream. We used the fast settle option of the processor for all recording channels. With this option, the recording of each channel was blanked out for the duration of each stimulation pulse on any electrode, and an additional 0.5 ms after. Electrical stimulation was controlled through a customized Python interface which allows easy configuration of the stimulation parameters and timing for each channel.

### 4.3 Data Collection and Task Description

We collected a dataset to train the models to predict the neural response and to find the best stimulation to induce a certain neural activity. Specifically, a set of stimulation parameters was used that has proven effective in inducing visual perception with intracortical electrodes, based on previous literature [17, 40], *i*.*e*., a train of 50 pulses lasting 167 ms (300 Hz), where each pulse is rectangular, cathodic first, and charge-balanced (cathodic and anodic duration of 170*µs*, interphase duration 60*µs*). For safety reasons, pulses within groups of simultaneously activated electrodes were temporally interleaved to minimize the instantaneous current delivered to the neural tissue (*i*.*e*., stimulating during the interpulse gap of other stimulating electrodes, Supplementary Figure 1C). For a stimulation train at 300 Hz and a single-pulse duration of 400 *µ*s there are 2.93 ms between pulses, during which pulses from different electrodes are distributed. We distributed up to six pulses from different electrodes; in case more than six electrodes were included in the stimulation pattern, up to 2 electrodes delivered pulses at the same time.

Three types of stimulation patterns were used:

1. **Random:** A random group of electrodes (between 1 and 10 total) was selected. For each selected electrode, a random stimulation current was chosen between 5 *µ*A to 50 *µ*A. Examples of random stimulation patterns are shown in Supplementary Figure 1A. N=5818
2. **Structured:** Occasionally, the patterns were not random but followed structured shapes such as rectangles, triangles, circles or lines. For these structured patterns the current from all the electrodes was set to 15*µA* or 45*µA*. Examples of structured stimulation patterns are shown in Supplementary Figure 1B. N=484.
3. **Single Electrode:** Additionally, we collected data to determine the current threshold and neural response for each individual electrode, which was then used as a single-electrode dataset to build the forward dictionary models. To find the minimum current needed to induce perception from each electrode, a binary search procedure was used [17]. This led to the collection of a dataset of neural responses to each single electrode stimulation for different currents. Note that this was not included in the training data for the forward and inverse neural network models. N=2389.

In the experiments, the participant was sitting comfortably in a chair, and a sequence of stimulation patterns was delivered to the UEA. After each stimulation, there was a minimum delay of 1 second before the next stimulus. Three tasks were performed, differing in the answers requested from the participant:

1. **No answers needed:** This allowed the collection of the neural response to a large number of stimulation patterns in a short amount of time. N=787.
2. **Detection Task:** The participant reported via keyboard whether the stimulation pattern induced a visual perception or not. N=4811 (25 % of which leading to perception).
3. **Description of the shape, brightness, color, and size of perception:** The participant reported the description by voice, and the experimenter collected the answer in an Excel table. Shape and size were free-form responses which did not vary much throughout data collection, brightness was rated between 0 and 5, and color was a free-form response, which was later grouped into 4 color categories (green, yellow, white, and ambiguous). This protocol took the longest amount of time, so fewer stimulation parameters were tested. N=704.

When using optimized stimulation patterns, task 2 was used, except the patterns of stimulation were no longer random, but determined by the output of the various stimulation synthesis methods.

### 4.4 Neural Activity Extraction

We calculated the neural activity for each stimulation using MUA_e_, an averaged representation of the aggregate spiking activity from several neurons near the electrode tip, as defined in [40]. The raw signal is first filtered between 0.5 and 9 kHz, full-wave rectified, low-pass filtered at 200 Hz, and down-sampled at 1 kHz. Such multi-unit activity approaches are computationally efficient to estimate online and have been show to provide an accurate representation of neural activity [59] as well as robust performance on decoding tasks [60].

As a measure of the neural response induced by each stimulation pattern, we used the average MUA_e_ from 100 ms to 200 ms after the end of stimulation, normalized by subtracting the average MUA_e_ in a baseline window from −110 ms to −10 ms before stimulation (ΔMUA_e_). This time window was chosen to be the closest to stimulation offset without any stimulation artifacts, even on the stimulating electrode.

### 4.5 Metrics

We evaluated our models using three complementary metrics. All models were trained using mean squared error (MSE) and additionally evaluated using *R*^2^ and Earth Mover’s Distance (EMD). Note that all metrics were computed on single-trial neural responses without averaging.

For a batch of *n* samples, the MSE between neural responses **y**_*i*_ and 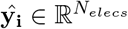 was defined as:

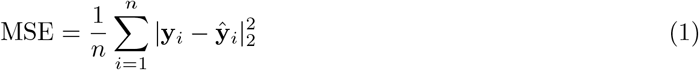

For forward models predicting neural responses, **y**_**i**_ was an observed neural response, and **ŷ**_**i**_ was a predicted response. For inverse models synthesizing new stimuli, **y**_**i**_ was the target neural response, and **ŷ**_**i**_ was an observed response to the stimulus synthesized for that target response.

The coefficient of determination (*R*^2^) measures explained variance for each channel. *R*^2^ was calculated for each channel individually and averaged across the array:

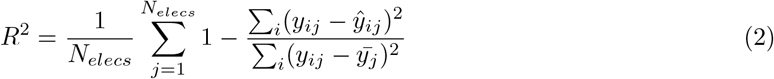

Here, *y*_*ij*_ and *ŷ*_*ij*_ represent the true and predicted neural response of the *j*th channel for the *i*th sample, and 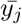 is the mean of all responses on the *j*th channel.

We also utilized Earth Mover’s Distance (EMD) to evaluate the proximity of two responses. EMD calculates the cost of “transporting” one sample into another, *i*.*e*., treating both neural responses as “piles of dirt”, and calculating the cost of moving the dirt from one response to the next. This metric accounts for the spatial characteristics of the electrode array–it considers responses that are “close” (e.g., off by one electrode) to have a small error–but is computationally expensive for use during training. EMD between two neural responses 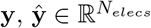 is defined as:

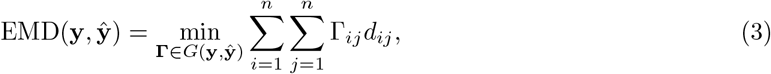

where

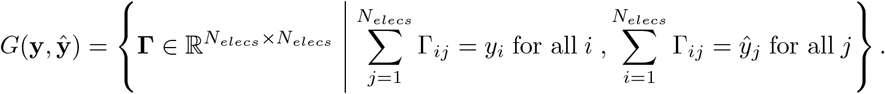

Here, *d*_*ij*_ is the physical distance between electrodes *i* and *j*, and *G*(**y, ŷ**) is the set of valid transport plans between two given recordings [48, 61]. This metric captures both the spatial relationship between electrodes and the relative magnitudes of neural responses.

### 4.6 Channel Inclusion Criteria

For the evaluation of forward and inverse models, we used only the channels in which any spiking activity was observed (Supplementary Figure 4A shows the electrodes with spiking activity resulting from spike sorting of 9 recordings lasting 5 minutes each in different days) and were ‘responsive’ to stimulation, meaning a variance in the neural response larger than 0.2 (Supplementary Figure 4B shows the variance in the neural response for each electrode). The total number of channels meeting these criteria is 54 out of 96 (Supplementary Figure 4C). These inclusion criteria ensure evaluating our models only in channels that were measuring neural activity.

### 4.7 Statistical Tests

Four different statistical tests were used to assess the significance of our results.

1. Wilcoxon signed-rank tests [62] were used in Section 2.1 to show that repeated stimulations had less variance within the same day than across days (Figure 2B) and that ΔMUA_e_ had less variance across repeated stimulations than post-stimulation MUA_e_ (Figure 2C).
2. A one-way ANOVA, with post-hoc independent Student’s *t* tests for individual differences, was used to test for differences between the forward neural network model and baselines (Section 2.1, Figure 2F). When controlling for multiple comparisons, we found that individual pairwise tests were not independent, prohibiting family-wise error rate based methods (e.g., Bonferroni), and thus instead corrected for multiple comparisons by controlling the false discovery rate [63].
3. A one-way repeated measures ANOVA was used to test for significant differences between stimulation synthesis methods (Section 2.2, Figure 3B, Figure 4B), with post-hoc paired Student’s *t* tests, correcting for multiple comparisons by controlling the false discovery rate [63].
4. A one-way ANOVA was used to test for differences across predictor models (Figure 6). For comparing each perception model’s performance with the stimulus-only baseline, paired Student’s *t* tests were used; p-values were corrected for multiple comparisons by controlling the false discovery rate [63].

### 4.8 Forward Model

We developed a deep neural network (‘Forward NN’) to predict the ΔMUA_e_ elicited from each stimulation pattern. The model is given as input both the stimulation pattern **s** (currents on each of 96 electrodes) and the average pre-stimulation activity (−110ms to 10ms) for each electrode on a given day 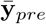, allowing it to account for daily variations in neural responses. The weights *θ*_1_ were trained to minimize the MSE loss:

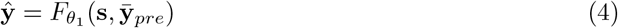

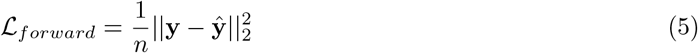

The architecture of the forward model consisted of 10 fully connected layers, with 20% dropout, batch normalization, and residual connections. The stimulation pattern was first input through 4 fully connected layers, with residual connections every 2 layers. This was then concatenated with the average pre-stimulation activity for the day and fed through 6 fully connected layers, with batch normalization and residual connections every 3 layers. ReLU activation functions were applied after each fully connected layer, with the final layer outputting 96 units corresponding to the predicted ΔMUA_e_ for each electrode. In total, the model had approximately 1.2 million trainable parameters.

The model was trained to minimize MSE between predicted and observed responses. The Adam optimizer was used with a learning rate of 0.0001 and a batch size of 256 for model training. A scheduler was used to decrease the learning rate by a factor of 0.5 every 200 epochs. Early stopping was also used to terminate training early based on a validation set performance, resulting in a total of 911 training epochs.

The ‘Forward NN’ was compared against several baselines:

#### Forward without pre-activity

To test the effect of ablating the average pre-stimulation activity, we trained an otherwise identical neural network to the forward model, but with only the stimulus as input.

#### 1-to-1

Each stimulating electrode was assumed to focally activate only the nearby neurons. Thus, the predicted response on each channel was directly proportional to the stimulating current on that electrode. The maximum observed neural activity was assumed to be proportional to the maximally delivered current (50*µA*)

#### Dictionary

Loosely inspired by [27], this method predicts multi-electrode stimulation responses via a lookup ‘dictionary’ of responses to single-electrode stimuli. Given an input multi-electrode stimulation 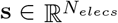 and selected channel *i ∈* [0, *N*_*elecs*_], then 𝒟*{***s**, *i}* is a dictionary function which outputs the average response in the training dataset across single-electrode stimuli on electrode *i* with the nearest amplitude to *s*_*i*_ (*e*.*g*. if *s*_2_ was 40*µA*, and the training set had 5 responses to single-electrode stimuli on electrode 2 at both 45 and 25*µA*, then 𝒟{**s**, 2} would return the average of the 5 45*µA* responses). We evaluated two dictionary variants. First, the neural responses for each individual single-electrode stimulus were averaged to get the predicted multi-electrode response (‘Dictionary’): 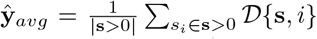; and second, the ideal linear combination of the single electrode responses was computed (‘Optimal Dictionary’): 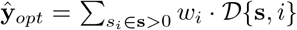 where *w*_*i*_ are learned weights optimized for each given stimulus and its true neural response **y**. This idealized dictionary would be infeasible to deploy in a real device, but gives a baseline of the maximum performance assuming linear combination of single-electrode responses.

#### Linear

A linear regression model was trained on the same dataset as the deep neural network. This model was given both the stimulus vector and the pre-activity to enable a fair comparison to the neural network forward model. The model can be formulated as: **ŷ**_*linear*_ = **Ws** + **b**, where **s** is the stimulus concatenated with the average pre-stimulation activity for that day, **W** is a weight matrix and **b** is a bias term.

#### Random

This approach randomly paired stimuli with neural activities from the training set, and averaged across 500 random pairings, serving as a lower performance bound.

### 4.9 Inverse Models

We developed several inversion methods using the forward model to target particular neural activity patterns. We compare our inversion methods with several baseline and state-of-the-art inversion methods.

#### Gradient

We developed a gradient-based optimization method, which freezes the training parameters of the forward model and uses back propagation to maximize the input stimulation pattern for a desired target neural response. This method directly optimizes a stimulation pattern **s** to minimize the difference between the forward model’s predicted response *F* (**s**) and the target response **y**, with L1 regularization on the stimulus to minimize stimulation amplitudes and the number of active electrodes:

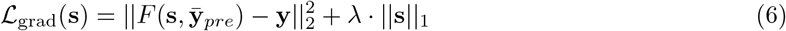

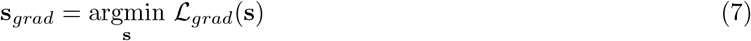

Where again 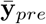 is the average pre-stimulation activity for the given day. We employed the Adam optimizer with learning rate of 0.001 and ran optimization for 10000 iterations or until convergence.

#### Inverse NN

We also built an inverse neural network model to directly map from stimuli to target neural responses. In this method, the weights of the forward model were frozen, and a secondary neural network was trained together with the frozen forward model in an autoencoder fashion, to approximate the forward model’s inverse function *F* ^*−*1^, with weights *θ*_2_. The architecture was the same as our forward model with three fully-connected layers of 96 units each, residual connections every other layer, and ReLU activations throughout the network, with dropout layers (rate=0.3) applied after each hidden layer to prevent overfitting. The inverse network also took as input the average pre-stimulation activity 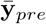 to account for daily variations in neural activity. The networks output was scaled by a factor *α* to ensure generated stimulation patterns fell within the valid range (0-50 *µ*A). The weights of the network *θ*_2_ were trained to minimize:

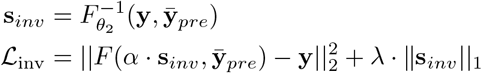

where *F* represents the frozen forward model, *F* ^*−*1^ is our inverse network, and **y** is the target neural response pattern. With this loss, the inverse network is trained to generate stimulation patterns that, when passed through the forward model, reproduce the target response.

#### 1 to 1 mapping

For our baseline comparisons we first implemented a conventional approach that assumes a direct relationship between stimulation and neural response. Here, stimulating amplitude was directly proportional to the desired activity on the underlying channel. Thus, there was a 1-1 mapping between target activations on each channel and stimulating currents on the same electrode. A target activity of 0 would have a current of 0, and a target activity of 4 (the maximum among our targets) would have a stimulating amplitude of 50*µA*

#### Dictionary

We evaluated an inverse dictionary approach, which leverages the dictionary (𝒟) of recorded neural responses 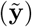 and stimulation pattern 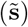 pairs to optimize stimulation patterns for a new target **y**. For a given target activation pattern, the dictionary stimulus was defined as the weighted average of every stimulus in the dictionary, weighted by the mean recorded response on channels active in the target response, similar to a traditional spike-triggered average.

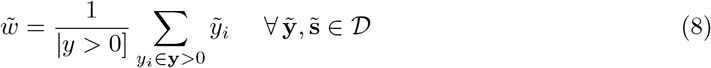

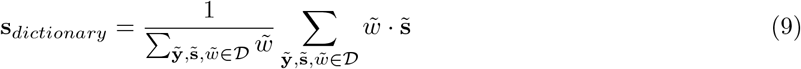

#### Linear

We also evaluated a linear regression model trained to map from neural responses back to stimulation patterns:

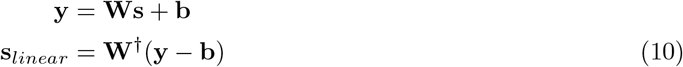

where *W* and *b* are learned parameters optimized to minimize MSE on the training data and *W* ^*†*^ is the pseudo inverse.

#### Replay Stimulus

We evaluated the activity generated by replaying the stimulation that evoked the target activity for Natural targets. This approach serves as an upper performance bound (the best achievable performance is limited by the variability of the response to the same stimulus).

#### Random

The performance of a random stimulation synthesis strategy for each target response was calculated using the average distance (measured using each metric) from each target to each response to random stimuli in the forward model training dataset.

For safety constraints, for all methods we only select the ten electrodes of **s** with the largest current amplitude.

### 4.10 Latent Factor Analysis

To quantify the manifold underlying neural activity we used latent factor analysis [54], which has been widely used previously (*e*.*g*. [55]). Under this model, neural responses 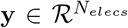 are assumed to be distributed according to a number of latent factors **z** *∈* ℛ^*D*^; **z** *∼* 𝒩(0, **I**):

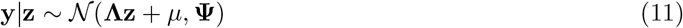

where 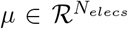 is the mean response for each channel, 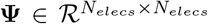 is a diagonal covariance matrix, and 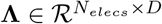 contains loadings mapping latent factors back to the neural activity.

**Λ** and **Ψ** were estimated using expectation-maximization with the python package scikit-learn [64], fit to the same dataset of neural responses used to fit the forward model. We chose *D* = 10, which captured 95% of the manifold variance in the observed responses (*i*.*e*., after accounting for noise). Similar to [55], the neural manifold was defined as the column space of **Λ**. The reported manifold distance was calculated as the Euclidean distance from each neural response to its orthogonal projection onto the columns of **Λ** (Equation 12). Note that even for the observed neural responses themselves, there is still a nonzero distance to the underlying manifold due to noise and variance not captured in the 10 components (Figure 4E, green dotted line). Natural targets had similar manifold distances as the neural manifold responses, and synthetic targets had significantly larger manifold distances.

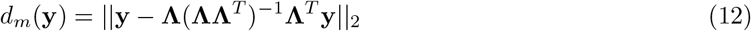

### 4.11 Predicting Perception

To validate the relevance of neural responses to perceptual measurements, we trained a decoder to predict verbal perceptual responses from recorded activity (descriptions of phosphene detection, color, and brightness rating). See Section 4.3 for more details on the task description.

The perception model used five fully connected layers of 256 units each with residual connections to every other layer; batch norm layers, dropout layers, and ReLU activations after each hidden layer. Training was performed using the Adam optimizer and cross-entropy loss. For each model type, dropout, batch size, ΔMUA_e_ window, dropout rates, and learning rate were tuned independently for each model configuration, and number of parameters was balanced across model types, to ensure fair comparison. Models were tested using leave-one-day-out cross validation to evaluate generalization across sessions.

The model receives as input neural data, either pre- and post-stimulation ΔMUA_e_, or the stimulation amplitudes per electrode. The model outputs were categorical variables corresponding to the perceptual attributes: binary classification for phosphene detection (seen/not seen), multi-class classification for color perception, and ordinal classification for brightness ratings.

## 5 Data and Code Availability

The data and code used for this study will be made available at https://github.com/jgranley/deep_neural_shaping

## 6 Acknowledgments

This work was financially supported by:

Ministerio de Ciencia, Innovación y Universidades Grant DTS19/00175.

Ministerio de Ciencia, Innovación y Universidades Grant PDC2022-133952-100.

European Union’s Horizon 2020 Research and Innovation Programme under Grant Agreement No. 899287 (NeuraViPeR).

European Union’s Horizon 2020 Research and Innovation Programme under the Marie Skłodowska-Curie grant agreement No 861423 (enTRAIN Vision).

Innovative Neurotechnology for Society (INTENSE). Dutch Neurotechnology Consortium.

Generalitat Valenciana, Directorate General of Science and Research, PROMETEO Grant CIPROM/2023/25.

Swiss National Science Foundation projects CA-DNNEdge (208227)

National Library of Medicine of the National Institutes of Health under Award Number DP2-LM014268 to MB. The content is solely the responsibility of the authors and does not necessarily represent the official views of the National Institutes of Health.

## 7 Author contributions

- **Conceptualization:** PM, JG, FG, MB, SL, and EF collaboratively designed the study and developed the core research questions, with critical input from LS, AL, CS, and RL.
- **Data collection & curation:** EF led participant recruitment, oversaw surgical procedures, and managed ethics approvals and clinical compliance. FG led experimental data collection and protocol development. LS contributed substantially to both data collection and quality assurance. CS supported data collection through participant oversight and in-room experimental supervision. PM, JG, AV, and RL contributed during select sessions.
- **Methods development:** JG and PM developed computational models, with input from FG. AV performed and validated spike sorting. All methods were reviewed by SL, MB, and EF.
- **Data analysis:** FG, PM, and JG conducted data analysis and visualization.
- **Writing:** FG, JG, and PM drafted the manuscript, prepared initial figures, and coordinated the revisions with critical input from AL, LS, CS, SL, MB, and EF. All authors approved the final version of the manuscript.
- **Project supervision & administration:** SL, MB, and EF jointly provided project oversight and scientific mentorship.

## 9 Supplementary Figures

**Supplementary Figure 1:**
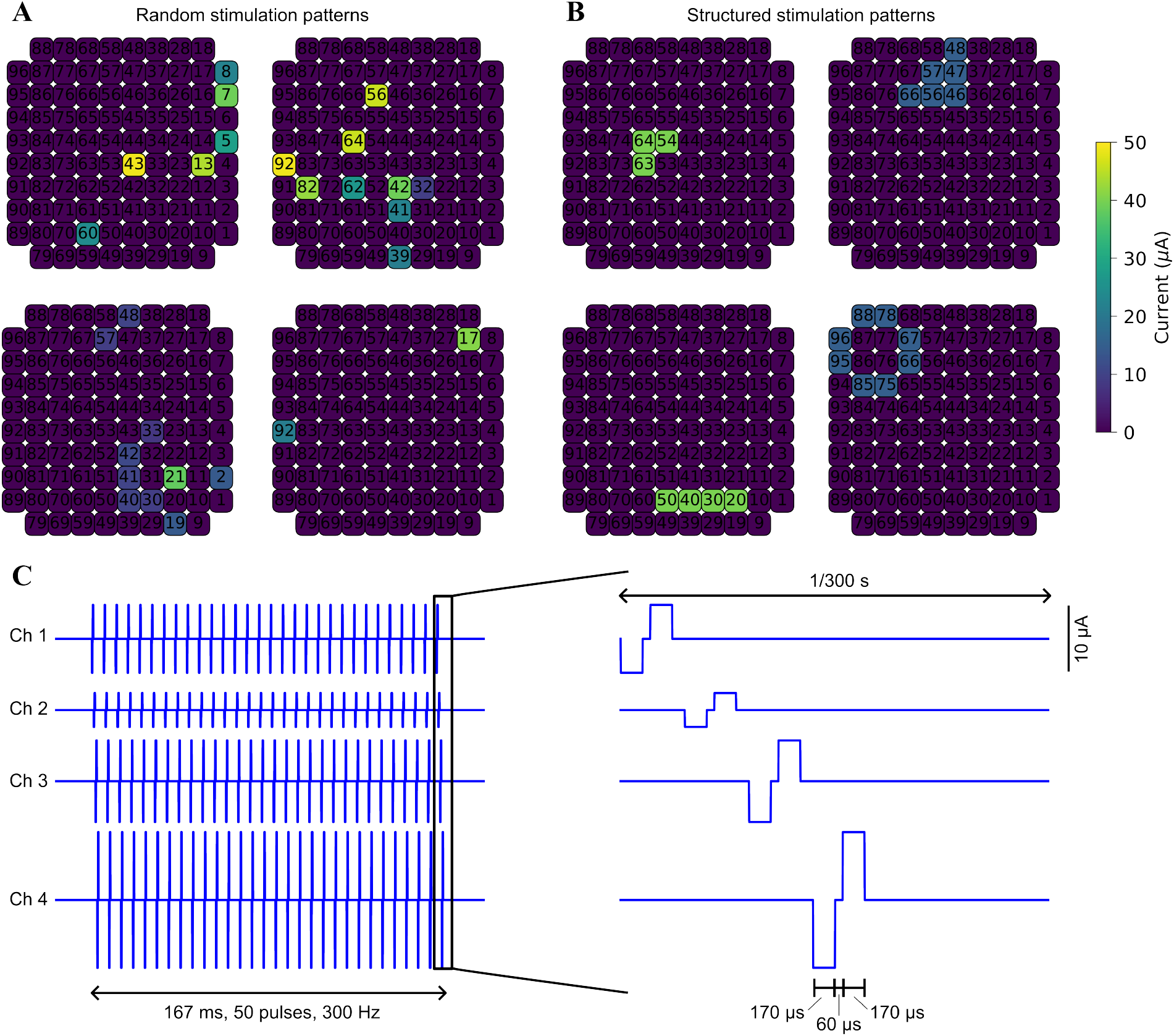
Stimulation patterns. A) Examples of random stimulation patterns. B) Examples of structured stimulation patterns. C) Example of a stimulation train from 4 electrodes (left) and zoom into the stimulation pulses (right).

**Supplementary Figure 2:**
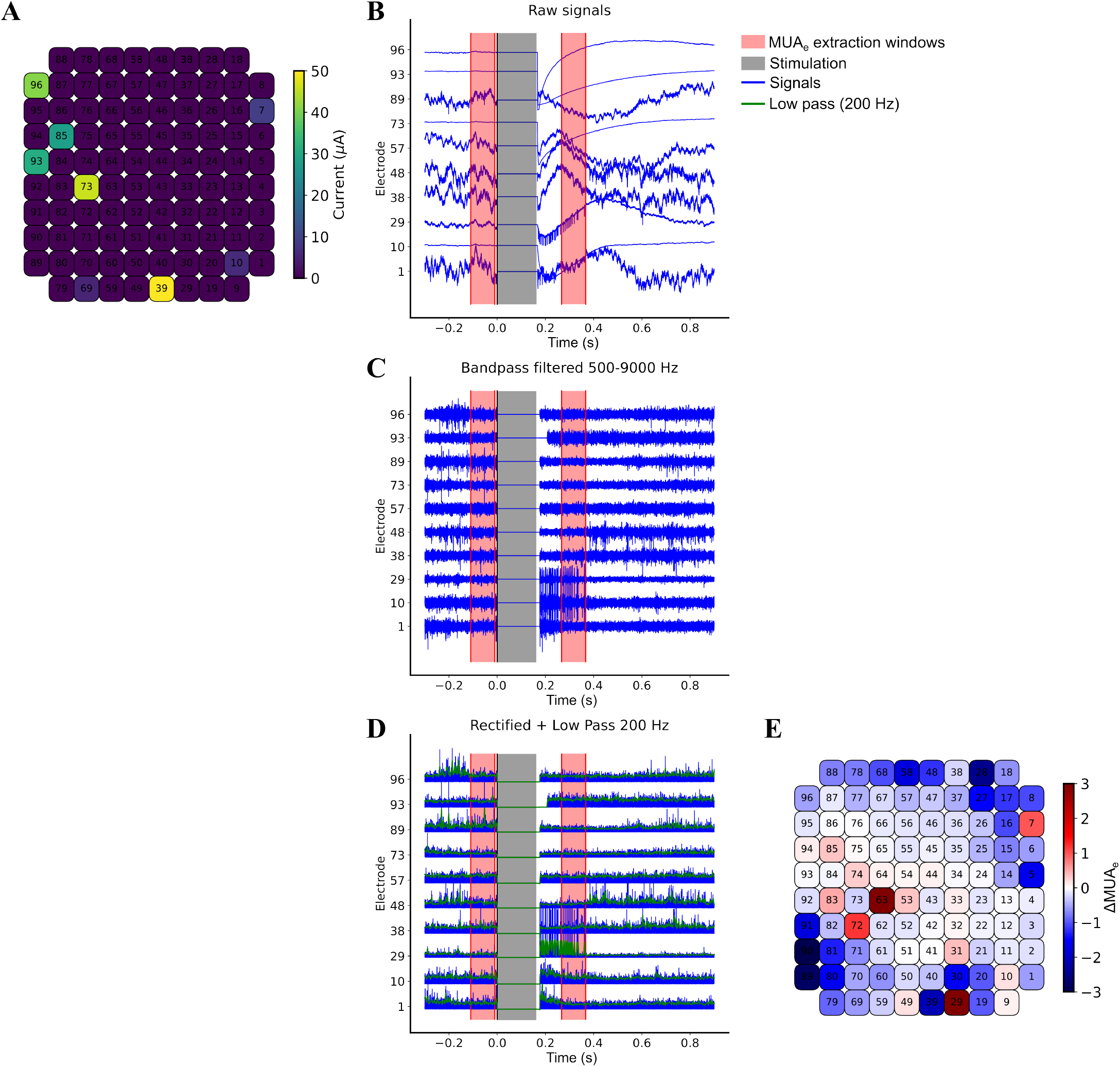
Extraction of the neural activity. A) Stimulation pattern. B) Raw signal. The black line represents the start of the stimulation, red lines represent the window used to extract the activity. C) Bandpass filtered signal (500-9000 Hz) D) Rectified signal (blue traces) and low pass filtered signal (200 Hz, green traces). E) Neural activity (ΔMUA_e_).

**Supplementary Figure 3:**
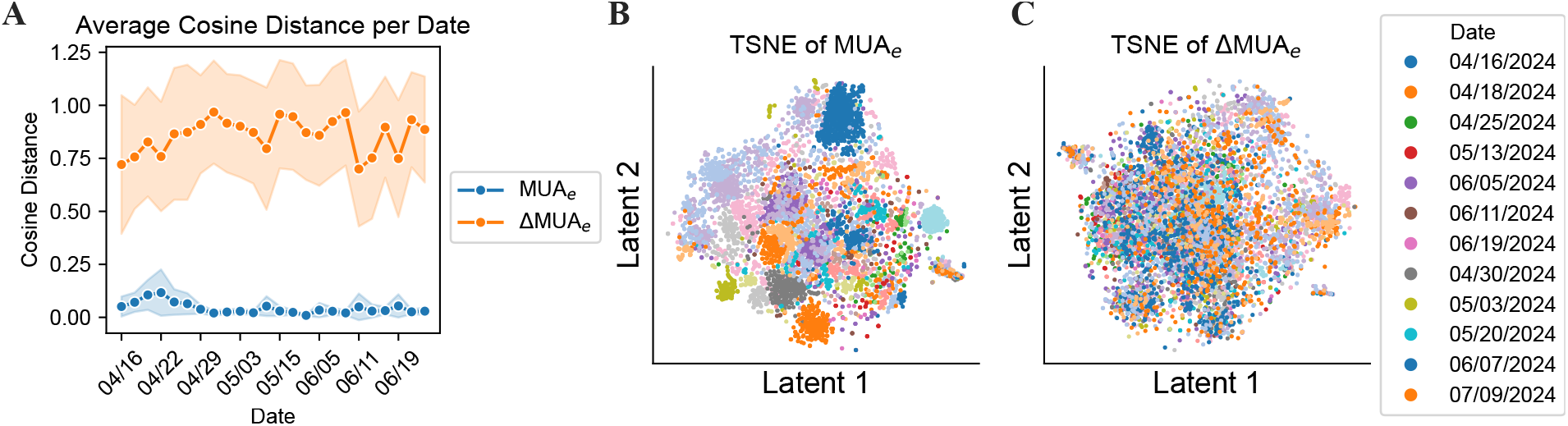
ΔMUA_e_ and post-stimulus MUA_e_ comparison of cosine distance (A) and t-distributed stochastic neighbor embedding (t-SNE) (B,C). We found that ΔMUA_e_ exhibited a higher cosine distance between points for a given day across all days, compared to the post-stimulation MUA_e_ (Supplementary Figure 3A). This suggests that the raw recordings contain a significant correlation structure dominated by a day-based baseline, which is mitigated in the ΔMUA_e_. Visualization of the data using t-distributed stochastic neighbor embedding (t-SNE) further supported this finding. The post-stimulation MUA_e_ showed much stronger day-wise clustering compared to ΔMUA_e_ (Supplementary Figure 3B,C). This indicates that the ΔMUA_e_ may be less dependent on the short-term neural state of a given day.

**Supplementary Figure 4:**
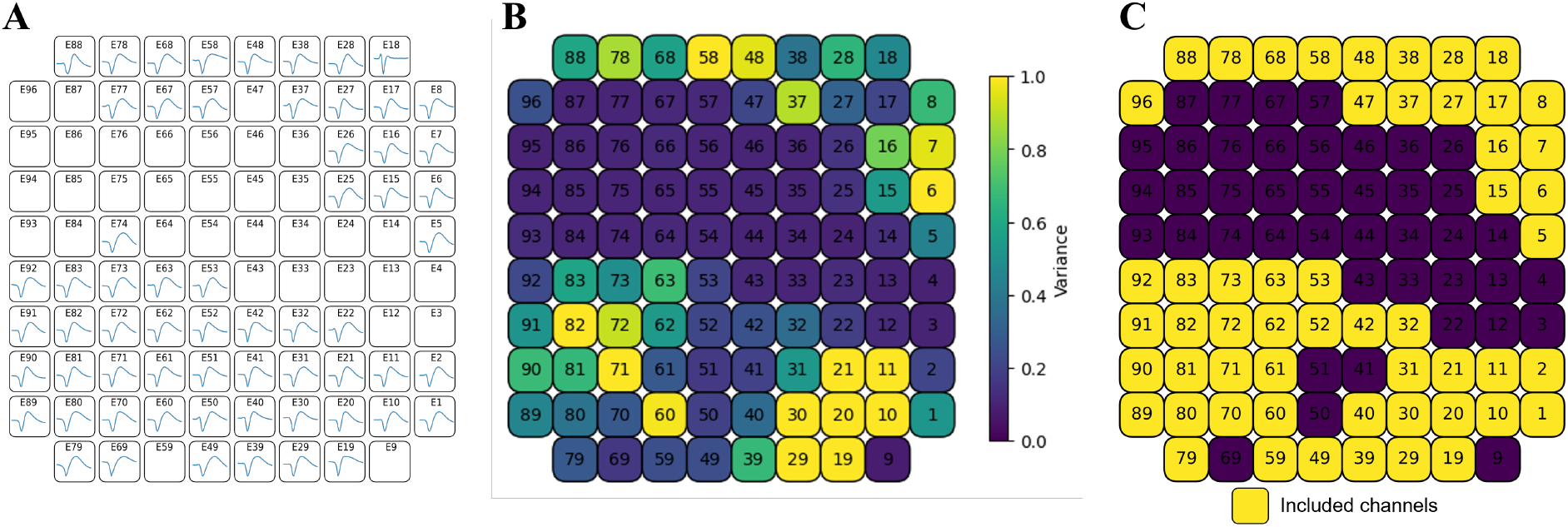
A) Spiking electrodes with manual spike sorting. B) Variance of the neural response to stimulation for each electrode. C) Included electrodes.

**Supplementary Figure 5:**
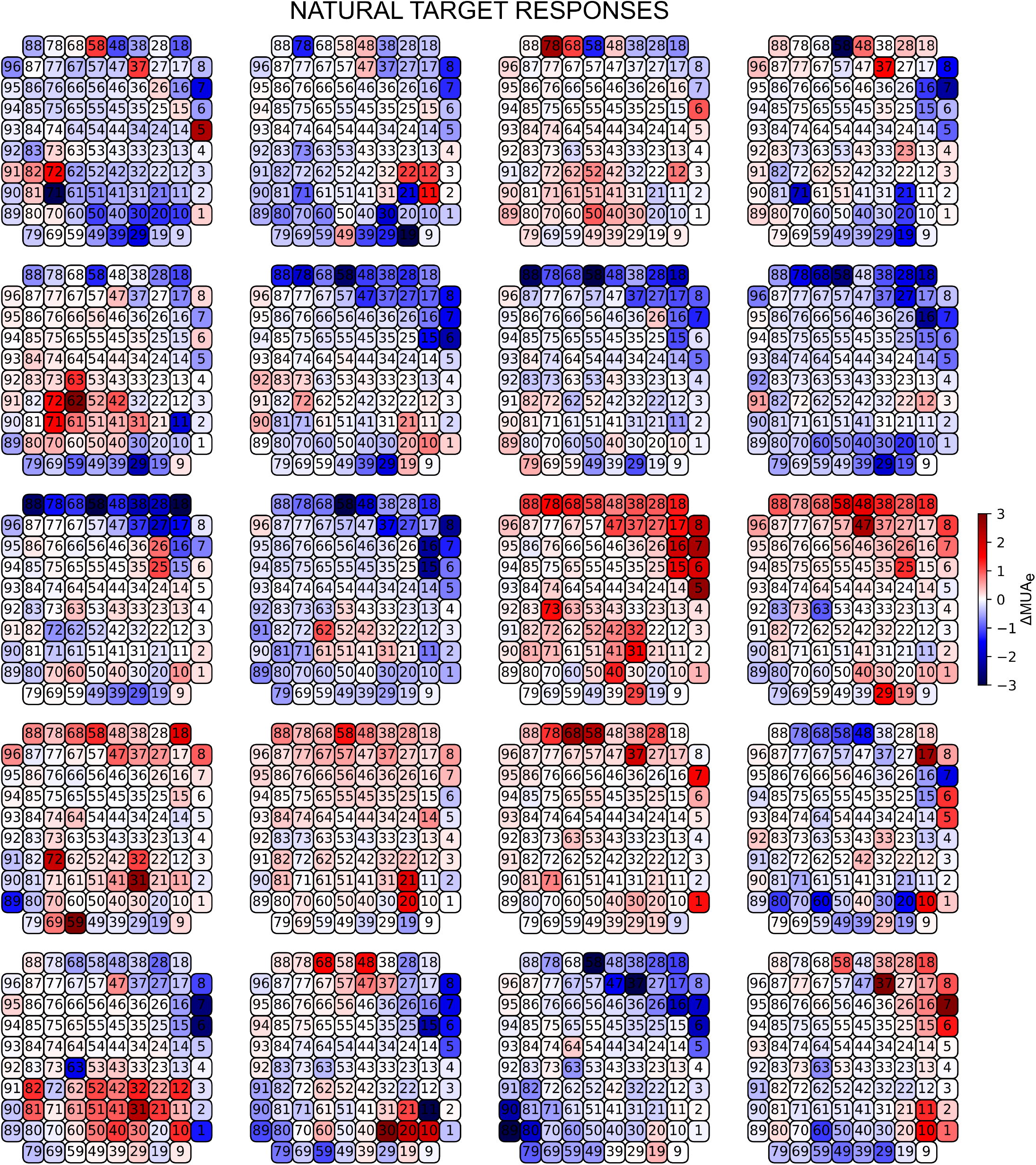
Natural Target Responses.

**Supplementary Figure 6:**
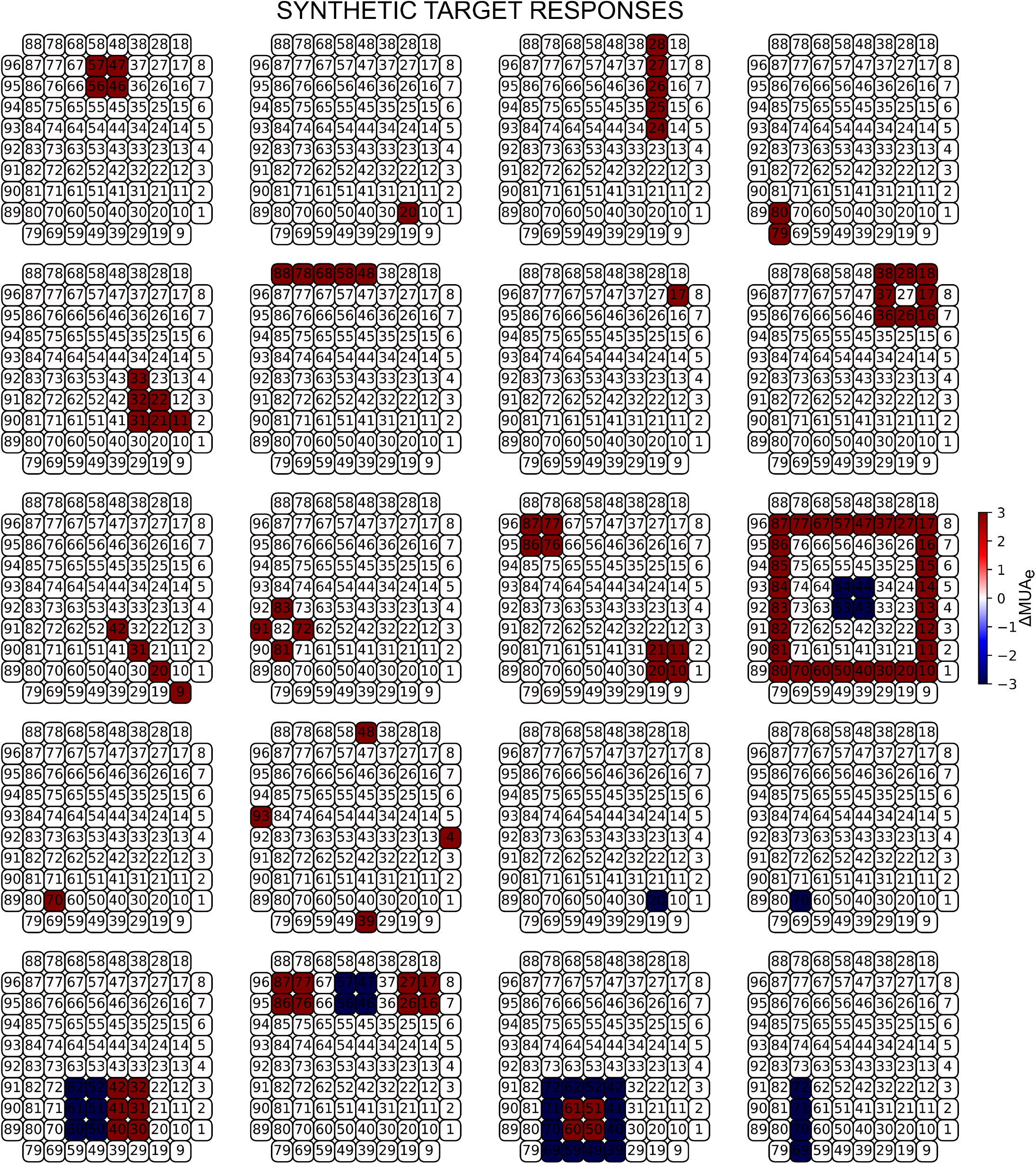
Synthetic Target Responses.

**Supplementary Figure 7:**
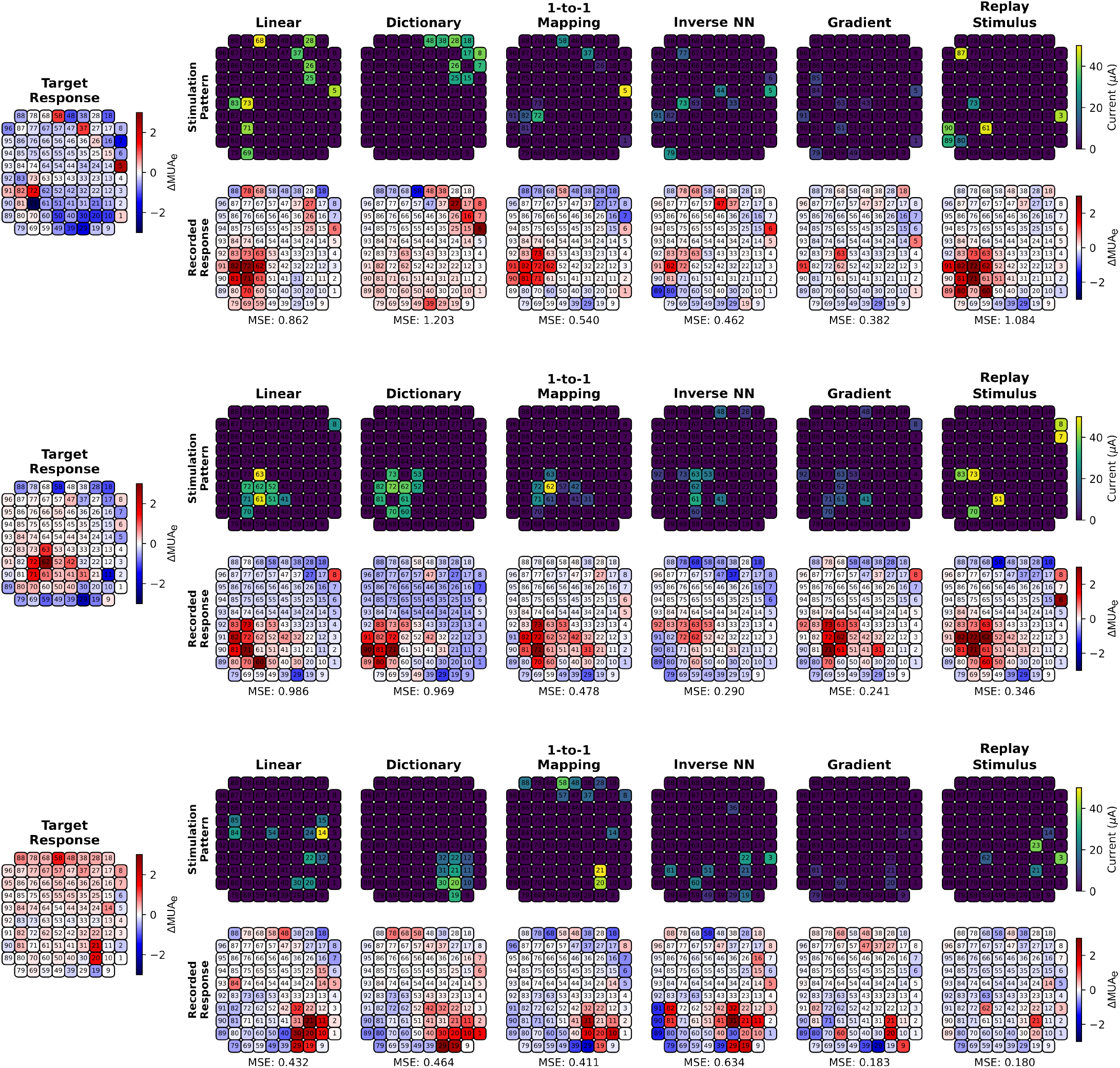
Natural Target Responses examples. For 3 examples of target activities (left): stimulation pattern proposed by each method (top) and neural response obtained (bottom).

**Supplementary Figure 8:**
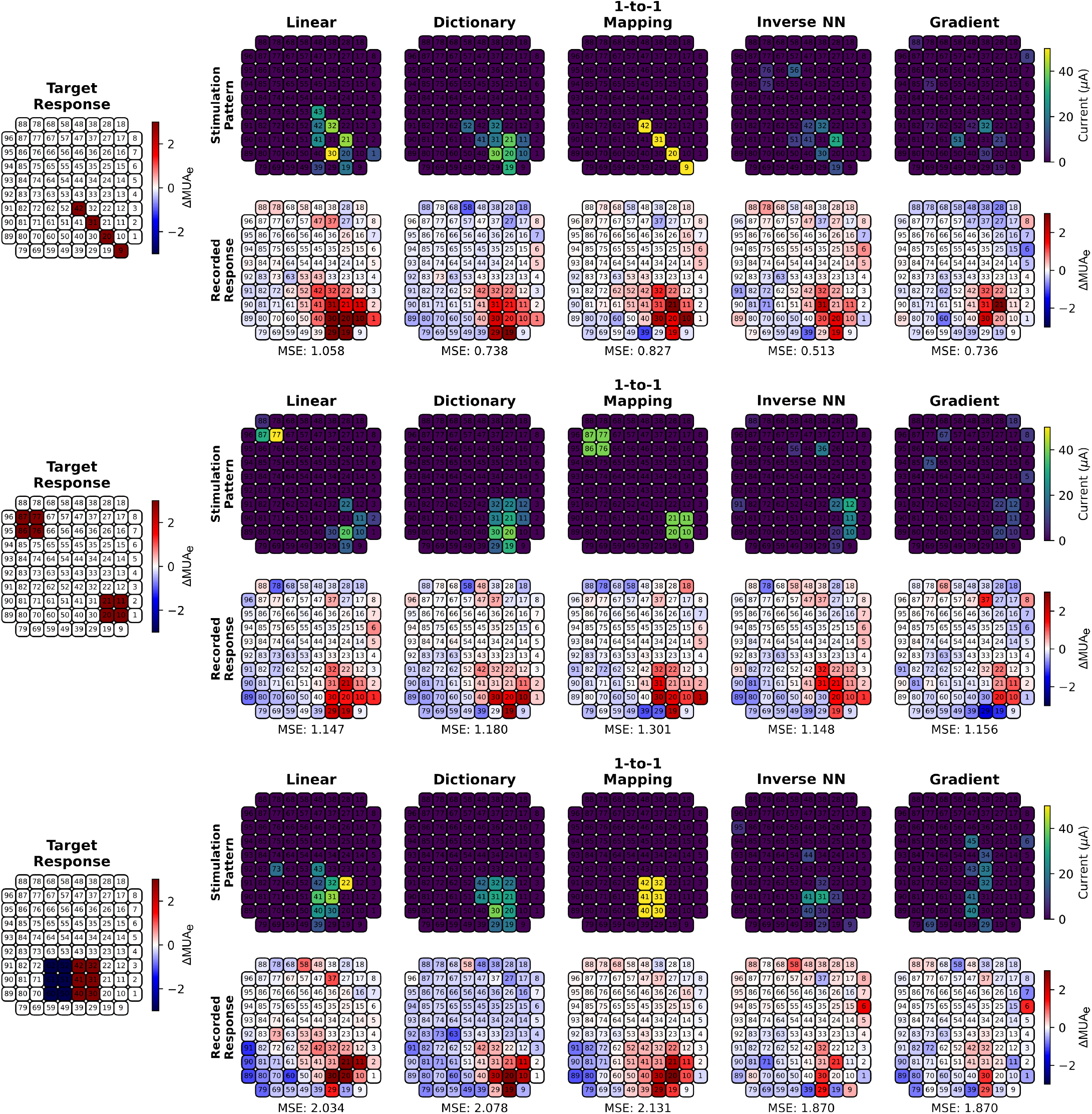
Synthetic Target Responses examples. For 3 examples of target activities (left): stimulation pattern proposed by each method (top) and neural response obtained (bottom).

## References

1. Lewis, P. M., Ackland, H. M., Lowery, A. J. & Rosenfeld, J. V. Restoration of vision in blind individuals using bionic devices: A review with a focus on cortical visual prostheses. Brain Research 1595, 51–73. ISSN: 0006-8993. https://www.sciencedirect.com/science/article/pii/S0006899314015674 (2024) (Jan. 2015).

2. Liu, X. et al. A narrative review of cortical visual prosthesis systems: the latest progress and significance of nanotechnology for the future. Annals of Translational Medicine 10, 716. ISSN: 2305-5839. https://www.ncbi.nlm.nih.gov/pmc/articles/PMC9279795/ (2024) (June 2022).

3. Pio-Lopez, L., Poulkouras, R. & Depannemaecker, D. Visual cortical prosthesis: an electrical perspective. Journal of Medical Engineering & Technology 45. Publisher: Taylor & Francis, 394–407. ISSN: 0309-1902. https://doi.org/10.1080/03091902.2021.1907468 (2024) (July 2021).

4. Zeng, F.-G., Rebscher, S., Harrison, W., Sun, X. & Feng, H. Cochlear Implants: System Design, Integration, and Evaluation. IEEE Reviews in Biomedical Engineering 1, 115–142. ISSN: 1941-1189. https://ieeexplore.ieee.org/abstract/document/4664429 (2024) (2008).

5. Flesher, S. N. et al. Intracortical microstimulation of human somatosensory cortex. en. Science Translational Medicine 8. ISSN: 1946-6234, 1946-6242. https://www.science.org/doi/10.1126/scitranslmed.aaf8083 (2024) (Oct. 2016).

6. Bjånes, D. A. & Moritz, C. T. A Robust Encoding Scheme for Delivering Artificial Sensory Information via Direct Brain Stimulation. IEEE Transactions on Neural Systems and Rehabilitation Engineering 27, 1994–2004. ISSN: 1558-0210. https://ieeexplore.ieee.org/abstract/document/8809212 (2024) (Oct. 2019).

7. Erickson-Davis, C. & Korzybska, H. What do blind people “see” with retinal prostheses? Observations and qualitative reports of epiretinal implant users. en. PLOS ONE 16. Publisher: Public Library of Science, e0229189. ISSN: 1932-6203. https://journals.plos.org/plosone/article?id=10.1371/journal.pone.0229189 (2024) (Feb. 2021).

8. Murray, C. D. en. in Psychoprosthetics (eds Gallagher, P., Desmond, D. & MacLachlan, M.) 119–129 (Springer, London, 2008). ISBN: 978-1-84628-980-4. https://doi.org/10.1007/978-1-84628-980-4_9 (2022).

9. Barry, M. P. et al. Video-mode percepts are smaller than sums of single-electrode phosphenes with the Orion® visual cortical prosthesis. Investigative Ophthalmology & Visual Science 61, 927. ISSN: 1552-5783 (June 2020).

10. Dagnelie, G. et al. Constructing a phosphene map for the inaugural recipient of the intracortical visual prosthesis (ICVP). Investigative Ophthalmology & Visual Science 64, 5520. ISSN: 1552-5783 (June 2023).

11. Luo, Y. H.-L. & da Cruz, L. The Argus® II Retinal Prosthesis System. Progress in Retinal and Eye Research 50, 89–107. ISSN: 1350-9462. https://www.sciencedirect.com/science/article/pii/S1350946215000701 (2024) (Jan. 2016).

12. Ayton, L. N. et al. First-in-Human Trial of a Novel Suprachoroidal Retinal Prosthesis. en. PLoS ONE 9 (ed Mori, K.) e115239. ISSN: 1932-6203. https://dx.plos.org/10.1371/journal.pone.0115239 (2024) (Dec. 2014).

13. Stingl, K. et al. Subretinal Visual Implant Alpha IMS – Clinical trial interim report. Vision Research. Sight restoration: prosthetics, optogenetics and gene therapy 111, 149–160. ISSN: 0042-6989. https://www.sciencedirect.com/science/article/pii/S0042698915000784 (2024) (June 2015).

14. Palanker, D., Le Mer, Y., Mohand-Said, S. & Sahel, J. A. Simultaneous perception of prosthetic and natural vision in AMD patients. en. Nature Communications 13. Publisher: Nature Publishing Group, 513. ISSN: 2041-1723. https://www.nature.com/articles/s41467-022-28125-x (2024) (Jan. 2022).

15. Veraart, C., Wanet-Defalque, M.-C., Gérard, B., Vanlierde, A. & Delbeke, J. Pattern Recognition with the Optic Nerve Visual Prosthesis. en. Artificial Organs 27, 996–1004. ISSN: 1525-1594. https://onlinelibrary.wiley.com/doi/abs/10.1046/j.1525-1594.2003.07305.x (2024) (2003).

16. Panetsos, F., Sanchez-Jimenez, A., Diaz-de Cerio, E. R., Diaz-Guemes, I. & Sanchez, F. M. Consistent Phosphenes Generated by Electrical Microstimulation of the Visual Thalamus. An Experimental Approach for Thalamic Visual Neuroprostheses. English. Frontiers in Neuroscience 5. Publisher: Frontiers. ISSN: 1662-453X. https://www.frontiersin.org/journals/neuroscience/articles/10.3389/fnins.2011.00084/full (2024) (July 2011).

17. Fernández, E. et al. Visual percepts evoked with an intracortical 96-channel microelectrode array inserted in human occipital cortex. en. Journal of Clinical Investigation 131, e151331. ISSN: 1558-8238. https://www.jci.org/articles/view/151331 (2024) (Dec. 2021).

18. Barry, M. P. et al. Preliminary visual function for the first human with the Intracortical Visual Prosthesis (ICVP). Investigative Ophthalmology & Visual Science 64, 2842. ISSN: 1552-5783 (June 2023).

19. Musk, E. & Neuralink. An Integrated Brain-Machine Interface Platform With Thousands of Channels. EN. Journal of Medical Internet Research 21. Publisher: JMIR Publications Inc., Toronto, Canada, e16194. https://www.jmir.org/2019/10/e16194 (2024) (Oct. 2019).

20. Beauchamp, M. S. et al. Dynamic Stimulation of Visual Cortex Produces Form Vision in Sighted and Blind Humans. Cell 181, 774–783.e5. ISSN: 0092-8674. https://www.sciencedirect.com/science/article/pii/S0092867420304967 (2024) (may 2020).

21. Maynard, E. M., Nordhausen, C. T. & Normann, R. A. The Utah Intracortical Electrode Array: A recording structure for potential brain-computer interfaces. Electroencephalography and Clinical Neurophysiology 102, 228–239. ISSN: 0013-4694. https://www.sciencedirect.com/science/article/pii/S0013469496951760 (2024) (Mar. 1997).

22. Youssef, D., Wittig, J. H., Jackson, S., Inati, S. K. & Zaghloul, K. A. Neuronal Spiking Responses to Direct Electrical Microstimulation in the Human Cortex. en. Journal of Neuroscience 43. Publisher: Society for Neuroscience Section: Research Articles, 4448–4460. ISSN: 0270-6474, 1529-2401. https://www.jneurosci.org/content/43/24/4448 (2025) (June 2023).

23. Kumaravelu, K. & Grill, W. M. Neural mechanisms of the temporal response of cortical neurons to intracortical microstimulation. Brain Stimulation 17, 365–381. ISSN: 1935-861X. https://www.sciencedirect.com/science/article/pii/S1935861X24000482 (2025) (Mar. 2024).

24. Histed, M. H., Bonin, V. & Reid, R. C. Direct activation of sparse, distributed populations of cortical neurons by electrical microstimulation. eng. Neuron 63, 508–522. ISSN: 1097-4199 (Aug. 2009).

25. Ghaffari, D. H., Chang, Y.-C., Mirzakhalili, E. & Weiland, J. D. Closed-loop Optimization of Retinal Ganglion Cell Responses to Epiretinal Stimulation: A Computational Study in 2021 10th International IEEE/EMBS Conference on Neural Engineering (NER) ISSN: 1948-3554 (May 2021), 597– 600. https://ieeexplore.ieee.org/document/9441437/?arnumber=9441437 (2024).

26. Shah, N. P. et al. Optimization of Electrical Stimulation for a High-Fidelity Artificial Retina in 2019 9th International IEEE/EMBS Conference on Neural Engineering (NER) ISSN: 1948-3554 (Mar. 2019), 714–718. https://ieeexplore.ieee.org/abstract/document/8716987 (2025).

27. Shah, N. P. et al. Precise control of neural activity using dynamically optimized electrical stimulation. eLife 13 (eds Beyeler, M., Gold, J. I. & Shivdasani, M.) Publisher: eLife Sciences Publications, Ltd, e83424. ISSN: 2050-084X. https://doi.org/10.7554/eLife.83424 (2025) (Nov. 2024).

28. Micera, S. et al. A computational model to design wide field-of-view optic nerve neuroprostheses Aug. 2023. https://www.researchsquare.com/article/rs-3218482/v1 (2024).

29. Vasireddy, P. K. et al. Efficient Modeling and Calibration of Multi-Electrode Stimuli for Epiretinal Implants in 2023 11th International IEEE/EMBS Conference on Neural Engineering (NER) ISSN: 1948-3554 (Apr. 2023), 1–4. https://ieeexplore.ieee.org/document/10123907 (2024).

30. Granley, J., Relic, L. & Beyeler, M. Hybrid Neural Autoencoders for Stimulus Encoding in Visual and Other Sensory Neuroprostheses. en. Advances in Neural Information Processing Systems 35, 22671–22685. https://proceedings.neurips.cc/paper_files/paper/2022/hash/8e9a6582caa59fda0302349702965171-Abstract-Conference.html (2024) (Dec. 2022).

31. Granley, J., Fauvel, T., Chalk, M. & Beyeler, M. Human-in-the-Loop Optimization for Deep Stimulus Encoding in Visual Prostheses. en. Advances in Neural Information Processing Systems 36, 79376–79398. https://proceedings.neurips.cc/paper_files/paper/2023/hash/fb06bc3abcece7b8725a8b83b8fa363Abstract-Conference.html (2025) (Dec. 2023).

32. Spencer, M. J., Kameneva, T., Grayden, D. B., Meffin, H. & Burkitt, A. N. Global activity shaping strategies for a retinal implant. en. Journal of Neural Engineering 16. Publisher: IOP Publishing, 026008. ISSN: 1741-2552. https://dx.doi.org/10.1088/1741-2552/aaf071 (2025) (Jan. 2019).

33. Castro, D., Grayden, D. B., Meffin, H. & Spencer, M. Neural Activity Shaping in Visual Prostheses with Deep Learning en. Dec. 2023. https://www.biorxiv.org/content/10.1101/2023.12.20.572123v1 (2024).

34. De Ruyter van Steveninck, J., Güçlü, U., van Wezel, R. & van Gerven, M. End-to-end optimization of prosthetic vision. Journal of Vision 22, 20. ISSN: 1534-7362. https://doi.org/10.1167/jov.22.2.20 (2024) (Feb. 2022).

35. Relic, L., Zhang, B.Tuan, Y.-L. & Beyeler, M. Deep Learning–Based Perceptual Stimulus Encoder for Bionic Vision in Proceedings of the Augmented Humans International Conference 2022 (Association for Computing Machinery, New York, NY, USA, Apr. 2022), 323–325. ISBN: 978-1-4503-9632-5. https://dl.acm.org/doi/10.1145/3519391.3524034 (2025).

36. Van Der Grinten, M. et al. Towards biologically plausible phosphene simulation for the differentiable optimization of visual cortical prostheses. en. eLife 13, e85812. ISSN: 2050-084X. https://elifesciences.org/articles/85812 (2025) (Feb. 2024).

37. Küçükoğlu, B. et al. End-to-end Learning of Safe Stimulation Parameters for Cortical Neuroprosthetic Vision en. Pages: 2025.01.23.634543 Section: New Results. Jan. 2025. https://www.biorxiv.org/content/10.1101/2025.01.23.634543v1 (2025).

38. Tafazoli, S. et al. Learning to control the brain through adaptive closed-loop patterned stimulation. en. Journal of Neural Engineering 17. Publisher: IOP Publishing, 056007. ISSN: 1741-2552. https://dx.doi.org/10.1088/1741-2552/abb860 (2025) (Oct. 2020).

39. Romeni, S., Zoccolan, D. & Micera, S. A machine learning framework to optimize optic nerve electrical stimulation for vision restoration. eng. Patterns (New York, N.Y.) 2, 100286. ISSN: 2666-3899 (July 2021).

40. Chen, X., Wang, F., Fernandez, E. & Roelfsema, P. R. Shape perception via a high-channel-count neuroprosthesis in monkey visual cortex. en. Science 370, 1191–1196. ISSN: 0036-8075, 1095-9203. https://www.science.org/doi/10.1126/science.abd7435 (2024) (Dec. 2020).

41. Papale, P., Luca, D. D. & Roelfsema, P. R. Deep generative networks reveal the tuning of neurons in IT and predict their influence on visual perception en. Pages: 2024.10.09.617382 Section: New Results. Oct. 2024. https://www.biorxiv.org/content/10.1101/2024.10.09.617382v1 (2025).

42. Grani, F. Improving cortical visual prostheses using closed-loop stimulation approaches PhD thesis (2024). http://dspace.umh.es/handle/11000/35610.

43. Grani, F. et al. ADVANTAGES OF BIDIRECTIONAL CORTICAL VISUAL PROSTHESES en. in IBRO Neuroscience Reports 15 (Oct. 2023), S948–S949. https://linkinghub.elsevier.com/retrieve/pii/S2667242123020651 (2024).

44. Allison-Walker, T., Hagan, M. A., Meikle, S. J., Price, N. S. C. & Wong, Y. T. Local field potential phase modulates the evoked response to electrical stimulation in visual cortex. en. Journal of Neural Engineering 22. Publisher: IOP Publishing, 016009. ISSN: 1741-2552. https://dx.doi.org/10.1088/1741-2552/ada828 (2025) (Jan. 2025).

45. Allison-Walker, T. J., Ann Hagan, M., Chiang Price, N. S. & Tat Wong, Y. Local field potential phase modulates neural responses to intracortical electrical stimulation in 2020 42nd Annual International Conference of the IEEE Engineering in Medicine & Biology Society (EMBC) ISSN: 2694-0604 (July 2020), 3521–3524. https://ieeexplore.ieee.org/abstract/document/9176186 (2025).

46. Dugué, L., Marque, P. & VanRullen, R. The Phase of Ongoing Oscillations Mediates the Causal Relation between Brain Excitation and Visual Perception. en. The Journal of Neuroscience 31, 11889–11893. ISSN: 0270-6474, 1529-2401. https://www.jneurosci.org/lookup/doi/10.1523/JNEUROSCI.1161-11.2011 (2025) (Aug. 2011).

47. Grani, F., Soto-Sánchez, C., Fimia, A. & Fernández, E. Toward a personalized closed-loop stimulation of the visual cortex: Advances and challenges. Frontiers in Cellular Neuroscience 16, 1034270. ISSN: 1662-5102. https://www.frontiersin.org/articles/10.3389/fncel.2022.1034270/full (2024) (Dec. 2022).

48. Leo, J., Ge, E. & Li, S. Wasserstein Distance in Deep Learning en. SSRN Scholarly Paper. Rochester, NY, Feb. 2023. https://papers.ssrn.com/abstract=4368733 (2025).

49. Minai, Y., Soldado-Magraner, J., Smith, M. A. & Yu, B. M. MiSO: Optimizing brain stimulation to create neural activity states. en. Advances in Neural Information Processing Systems 37, 24126–24149. https://proceedings.neurips.cc/paper_files/paper/2024/hash/2af641762dc02035c31a9314b2d090b6-Abstract-Conference.html (2025) (Dec. 2024).

50. Bashivan, P., Kar, K. & DiCarlo, J. J. Neural population control via deep image synthesis. en. Science 364, eaav9436. ISSN: 0036-8075, 1095-9203. https://www.science.org/doi/10.1126/science.aav9436 (2024) (May 2019).

51. Walker, E. Y. et al. Inception loops discover what excites neurons most using deep predictive models. en. Nature Neuroscience 22. Publisher: Nature Publishing Group, 2060–2065. ISSN: 1546-1726. https://www.nature.com/articles/s41593-019-0517-x (2025) (Dec. 2019).

52. Morales-Gregorio, A. et al. Neural manifolds in V1 change with top-down signals from V4 targeting the foveal region. English. Cell Reports 43. Publisher: Elsevier. ISSN: 2211-1247. https://www.cell.com/cell-reports/abstract/S2211-1247(24)00699-5 (2025) (July 2024).

53. Langdon, C., Genkin, M. & Engel, T. A. A unifying perspective on neural manifolds and circuits for cognition. en. Nature Reviews Neuroscience 24. Publisher: Nature Publishing Group, 363–377. ISSN: 1471-0048. https://www.nature.com/articles/s41583-023-00693-x (2025) (June 2023).

54. Bartholomew, D. J., Knott, M. & Moustaki, I. Latent Variable Models and Factor Analysis: A Unified Approach en. Google-Books-ID: 5YxvE1S56DwC. ISBN: 978-1-119-97370-6 (John Wiley & Sons, June 2011).

55. Sadtler, P. T. et al. Neural constraints on learning. en. Nature 512. Publisher: Nature Publishing Group, 423–426. ISSN: 1476-4687. https://www.nature.com/articles/nature13665 (2025) (Aug.2014).

56. Stringer, C., Pachitariu, M., Steinmetz, N., Carandini, M. & Harris, K. D. High-dimensional geometry of population responses in visual cortex. en. Nature 571. Publisher: Nature Publishing Group, 361–365. ISSN: 1476-4687. https://www.nature.com/articles/s41586-019-1346-5 (2025) (July 2019).

57. Normann, R. A. & Fernandez, E. Clinical applications of penetrating neural interfaces and Utah Electrode Array technologies. Journal of Neural Engineering 13, 061003. ISSN: 1741-2560, 1741-2552. https://iopscience.iop.org/article/10.1088/1741-2560/13/6/061003 (2024) (Dec.2016).

58. Rocca, A. et al. Robot-assisted implantation of a microelectrode array in the occipital lobe as a visual prosthesis: technical note. Journal of Neurosurgery 140, 1169–1176. ISSN: 0022-3085, 1933-0693. https://thejns.org/view/journals/j-neurosurg/140/4/article-p1169.xml (2024)(Apr. 2024).

59. Trautmann, E. M. et al. Accurate Estimation of Neural Population Dynamics without Spike Sorting. English. Neuron 103. Publisher: Elsevier, 292–308.e4. ISSN: 0896-6273. https://www.cell.com/neuron/abstract/S0896-6273(19)30428-3 (2025) (July 2019).

60. Ahmadi, N., Constandinou, T. G. & Bouganis, C.-S. Robust and accurate decoding of hand kinematics from entire spiking activity using deep learning. en. Journal of Neural Engineering 18. Publisher: IOP Publishing, 026011. ISSN: 1741-2552. https://dx.doi.org/10.1088/1741-2552/abde8a (2025) (Feb. 2021).

61. Aoun, A., Shetler, O., Raghuraman, R., Rodriguez, G. A. & Hussaini, S. A. Beyond correlation: optimal transport metrics for characterizing representational stability and remapping in neurons encoding spatial memory. English. Frontiers in Cellular Neuroscience 17. Publisher: Frontiers. ISSN: 1662-5102. https://www.frontiersin.org/journals/cellular-neuroscience/articles/10.3389/fncel.2023.1273283/full (2025) (Jan. 2024).

62. Rosner, B., Glynn, R. J. & Lee, M.-L. T. The Wilcoxon signed rank test for paired comparisons of clustered data. eng. Biometrics 62, 185–192. ISSN: 0006-341X (Mar. 2006).

63. Benjamini, Y. & Yekutieli, D. The control of the false discovery rate in multiple testing under dependency. The Annals of Statistics 29. Publisher: Institute of Mathematical Statistics, 1165–1188. ISSN: 0090-5364, 2168-8966. https://projecteuclid.org/journals/annals-of-statistics/volume-29/issue-4/The-control-of-the-false-discovery-rate-in-multiple-testing/10.1214/aos/1013699998.full (2025) (Aug. 2001).

64. Pedregosa, F. et al. Scikit-learn: Machine Learning in Python. Journal of Machine Learning Research 12, 2825–2830 (2011).

